# Single-neuron models linking electrophysiology, morphology and transcriptomics across cortical cell types

**DOI:** 10.1101/2020.04.09.030239

**Authors:** Anirban Nandi, Tom Chartrand, Werner Van Geit, Anatoly Buchin, Zizhen Yao, Soo Yeun Lee, Yina Wei, Brian Kalmbach, Brian Lee, Ed Lein, Jim Berg, Uygar Sümbül, Christof Koch, Bosiljka Tasic, Costas A. Anastassiou

## Abstract

Identifying the cell types constituting brain circuits is a fundamental question in neuroscience and motivates the generation of taxonomies based on electrophysiological, morphological and molecular single cell properties. Establishing the correspondence across data modalities and understanding the underlying principles has proven challenging. Bio-realistic computational models offer the ability to probe cause-and-effect and have historically been used to explore phenomena at the single-neuron level. Here we introduce a computational optimization workflow used for the generation and evaluation of more than 130 million single neuron models with active conductances. These models were based on 230 *in vitro* electrophysiological experiments followed by morphological reconstruction from the mouse visual cortex. We show that distinct ion channel conductance vectors exist that distinguish between major cortical classes with passive and h-channel conductances emerging as particularly important for classification. Next, using models of genetically defined classes, we show that differences in specific conductances predicted from the models reflect differences in gene expression in excitatory and inhibitory cell types as experimentally validated by single-cell RNA-sequencing. The differences in these conductances, in turn, explain many of the electrophysiological differences observed between cell types. Finally, we show the robustness of the herein generated single-cell models as representations and realizations of specific cell types in face of biological variability and optimization complexity. Our computational effort generated models that reconcile major single-cell data modalities that define cell types allowing for causal relationships to be examined.

**Highlights:**

1. Generation and evaluation of more than 130 million single-cell models with active conductances along the reconstructed morphology faithfully recapitulate the electrophysiology of 230 *in vitro* experiments.
2. Optimized ion channel conductances along the cellular morphology (‘all-active’) are characteristic of model complexity and offer enhanced biophysical realism.
3. Ion channel conductance vectors of all-active models classify transcriptomically defined cell-types.
4. Cell type differences in ion channel conductances predicted by the models correlate with experimentally measured single-cell gene expression differences in inhibitory (Pvalb, Sst, Htr3a) and excitatory (Nr5a1, Rbp4) classes.
5. A set of ion channel conductances identified by comparing between cell type model populations explain electrophysiology differences between these types in simulations and brain slice experiments.
6. All-active models recapitulate multimodal properties of excitatory and inhibitory cell types offering a systematic and causal way of linking differences between them.

## 1 Introduction

The nervous system consists of cell classes or types that can be defined by specific molecular signatures^1,2^, morphologies ^3,4^ and electrophysiological properties ^5–8^. In recent years, extensive single-cell characterization of neurons, mainly propelled by advances in sequencing technology ^9^, is revealing a multitude of ‘cell types’ ^10–12^. Despite our ever-increasing ability to detect distinguishing molecular, morphological and electrophysiological features to differentiate between such types, unraveling causal relationships between data modalities has been difficult. For example, how does, a particular distribution of ion channel conductances dictated by the genome manifest itself in the various electrophysiological features recorded in slice experiments? Experimentally, this process would involve elaborate genetic and/or pharmacological manipulations that are difficult to scale.

Single-cell models can link various types of data by incorporating constraints and generating predictions across data modalities, e.g., predicting a particular ion channel conductance based on properties of the electrophysiological response such as spike shape or frequency. In particular, two types of models have attracted attention: i) models linking transcriptomic data (e.g. single-cell RNA sequencing) with electrophysiological properties (e.g. recorded via whole-cell patch clamping), and ii) models linking morphological data (e.g. dendritic reconstructions) with electrophysiological experiments. Regarding the former, a series of studies have successfully associated aspects of single-cell molecular and electrophysiological features in models ^7^, supporting and extending the links found experimentally ^16–19^. However, these studies also demonstrate significant cell-to-cell variability across data modalities, challenging the idea that a single model can faithfully capture any particular cell type. Regarding the latter type of models, numerous studies have used them to explore experimental observables such as action potential generation and backpropagation, dendritic Ca-events, etc. along the detailed cellular morphology of cortical neurons in rodent ^20–22^, non-human primate ^23^ and human ^24,25^. While valuable, a limitation of these models (i.e. combining electrophysiology and detailed morphology) is the vast parameter space rendering their generation difficult and computationally expensive. As a result, most studies have either focused on a limited number of experiments (ranging from a single cell to only a handful) or, alternatively, models offering reduced bio-realism, e.g., by neglecting conductances along the dendritic morphology ^26^. It follows that *en masse* model generation from a large number of single-cell experiments has hitherto proven difficult. This is particularly true for models that link all three key experimental modalities: single-cell electrophysiology, detailed morphology and molecular signatures.

Recently, two independently collected datasets shed light into the cortical cell type taxonomy. The first data defines transcriptomic cell types based on single-cell RNA sequencing ^1,2^. The second set uses the somatic electrophysiological responses of single neurons and morphology reconstructions from the same cell to classify cortical cell types ^8^. First, we developed a systematic model generation workflow that uses these electrophysiology and fully reconstructed morphology data to generate biologically faithful single-cell representations for each experiment for more than 230 excitatory and inhibitory single cells. This large-scale optimization effort resulted in a set of ion channel conductance values for each section of the single cell morphology (soma, axon initial segment, apical and basal dendrites), i.e., a reduced latent representation for each single-cell. Recapitulating biology ^27,28^, we generate models containing active (i.e. Na^+^, K^+^ and Ca^++^-dependent) conductances along the entirety of their somatic, dendritic and axonal sections resulting in ‘all-active’ conductance-based single-cell representations. Then, in a second step, we compared this latent representation of single cells based on their electrophysiology and morphology to corresponding gene expression profiles measured by single-cell transcriptomics.

We use these all-active models to address three important questions. First, do ion channel conductances distributed along the single-cell morphology capture the distinguishing characteristics and variability of major cortical cell classes? Second, can these conductance vectors explain distinguishing properties between cell types (e.g., can differential expression of specific ion channel genes explain differences at the level of electrophysiological properties these channels are known to impact)? Third, how unique are these single-cell model representations and to what extent do they represent realizations of a particular cell type? To address these questions, we evaluated more than 130 million all-active single-cell models to show that accounting for ion channels along the entire morphology results in markedly different and more realistic ion channel conductance vectors compared to simpler model representations. We also show that, despite considerable biological variability, ion conductance profiles distributed along various regions of the cellular morphology distinguish between major cortical classes. For a subset of excitatory and inhibitory cell classes, we show how pairwise comparison between all-active conductance vectors trained solely on electrophysiology and morphology data results in set of conductances corresponding to genes that are indeed shown to be differentially expressed by single-cell RNA-sequencing. In such manner, we are able to link the differentially expressed genes to distinct, cell type-specific electrophysiology properties. Finally, we show that biological variability, while unequivocally entering into ion channel conductance estimations, is not enough to abolish cell type-identity, and that within-experiment variability remains consistently smaller than within-class. We conclude that the all-active models link single-cell electrophysiology, morphology and transcriptomic properties offering a systematic and causal way of linking evidence across these data modalities.

## 2 Results

### 2.1 Large-scale generation of single-cell models

The excitatory and inhibitory neurons assayed in the current work originate from mouse primary visual cortex (V1) as well as neighboring cortical areas across all six cortical layers (Figure 1a). In our work, we use two independently collected data sets. The first set uses single-cell RNA sequencing to reveal the transcriptomic profiles of individual cells as gene expression counts over million reads ^1,2^. The second set ^8^ consists of somatic electrophysiological responses of single neurons to standardized current clamp protocols, as well as the morphologies of those same cells reconstructed via a semi-automated approach ^29^. Notably, transgenic Cre lines crossed to Cre-reporter lines were used in both datasets to genetically target specific groups of cells ^30^. Each of the 230 single-cell experiments considered here is characterized according to the cortical layer its soma is located (L1-6), its dendrite morphology type (‘spiny’ and ‘aspiny or sparsely spiny’, representative of putative excitatory and inhibitory type or the class of a neuron, respectively) as well as the corresponding Cre-line with 19 lines grouped into 4 ‘broad’ subclasses: pyramidal subclass represented by all excitatory Cre-lines together (Pyr; see **STAR Methods**) and three inhibitory subclasses expressing parvalbumin (Pvalb), somatostatin (Sst), or serotonin receptor 5Htr3a (Htr3a) ^26^ (Figure 1a-b). Throughout this manuscript we use the nomenclature introduced above, i.e. cell classes characterize the excitatory and inhibitory divide based on the morphology type and any refinement on these two classes are denoted as a subclass.

**Figure 1:**
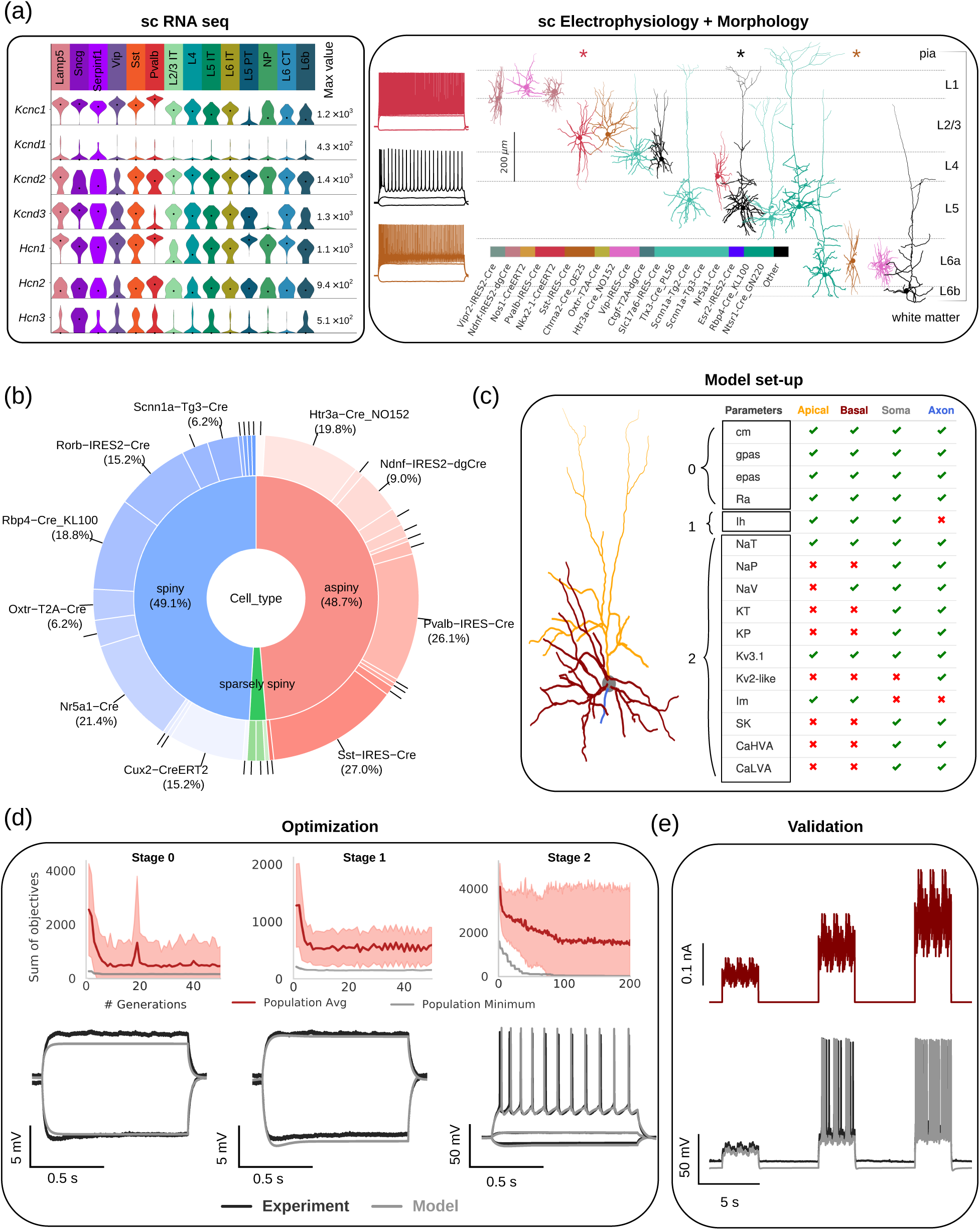
Cell types in mouse Visual Cortex (VIS) and single-cell model generation. **(a)** Data modalities used in the paper: (left) Single neuron RNA-sequencing (T) data ^1^ and (right) single neuron morpho-electric (ME) data ^8^. Sample morphologies arranged according to normalized depth from pia including putative cortical layer markings and electrophysiological recordings for 3 cells in the same set under subthreshold and suprathreshold current injections (colors: Cre-line). **(b)** Overview of the 230 modeled cells in the ME dataset based on dendrite type and Cre-line (cell class – spiny: putative excitatory, sparsely spiny and aspiny: putative inhibitory). **(c)** Model set-up: Morphology (CellID: 483101699) and predefined set of active conductances/passive properties marked according to their inclusion in each of the morphology sections (apical, basal dendrites, soma, AIS) **(d)** The optimization workflow is divided into 3 stages (Stage 0, 1, 2). The parameters added to the variable list at each stage are highlighted at the left of the table in **(c)**. top: Comparison between experimental traces (black) and the fitted model (grey) at the conclusion of each stage for a representative model (same CellID as in **(c)**). (bottom) Evolution of the sum of objectives as a function of generation number for the same model in (c). The best performing individual model (grey) at each generation and the average performance of all the individual models (red) of that generation (spread: standard deviation of the population). **(e)** Comparison between model and experiment spiking responses (bottom) under specific test stimulus (i.e., standardized colored noise (top)) to evaluate explained variance (see **STAR Methods**).

To generate biophysically realistic single-cell models, our computational workflow relied on two data modalities: the somatic electrophysiology response from whole-cell current-clamp experiments and the reconstructed morphology of each neuron. For a particular cell, the main challenge in the model generation is distributing a set of voltage-gated ion channel conductances along the morphology (Figure 1c) in order for the resulting model to replicate the experimental electrophysiology features. The passive properties of the membrane together with the maximal conductances (ḡ) for these voltage-gated Na^+^, K^+^ and Ca^++^-channels can be represented with a vector which uniquely identifies every model. Fitting the various conductances involves formulating a multi-objective optimization ^31^ problem, where each objective is a measure of the agreement between the experimental and model feature for the specific current input (dc steps) ^32,33^. The optimization leverages evolutionary algorithms ^34,35^ that iteratively search the space to improve upon previously evaluated solutions. We developed an automated model generation workflow that divides the fitting procedure in three stages: passive parameter fit (Stage 0), h-channel conductance density fit (Stage 1) and all-active parameter fit (Stage 2, Figure 1c; see also **STAR Methods** for the rationale behind adopting this workflow). The multi-objective optimization aims to reduce each objective independently (Figure 1d) resulting in the fit of a set of electrophysiology features (**Table S2,3**). In a final step, to validate the all-active model, we evaluated it on novel current stimuli (noise stimuli) that models were not trained on and compared features against experimental data from the same noise stimuli (Figure 1e).

To derive the single neuron models, we employed 15 models of voltage-gated ion channels (**Table S1**) known to be expressed in rodent cortical neurons (see **STAR Methods**, ^20,26^). During the optimization procedure, we kept the kinetics of these conductances unaltered and only allowed the maximal conductance density and the passive properties to vary. This approach resulted in a maximum of 43 variables per model along four identified regions of the morphology (soma, axon initial segment or AIS, basal and apical dendrites). To optimize these variables for a single all-active model, the workflow involved 300 generations (Stage 0: 50, Stage 1: 50, Stage 2: 200) with 512 new models generated and evaluated at each generation. The final model was selected from a group of 10 best models generated in this evolutionary procedure (termed “hall of fame”) for a single seed. Four seeds were used to randomize initialization from the same parameter bounds. In total, a single all-active model required the generation and evaluation of more than 600,000 models. Therefore, model generation for the entirety of the single-cell experiments required evaluation of more than 130 million single-cell models and 3.5 million CPU core-hours.

How well do the all-active models capture experimental features? To quantify the differences between model and experiment for any feature at a specified stimulus, we calculated the z-score for each feature in the best model resulting from the optimization process (see **STAR Methods**). Evaluating the error for the best model (i.e. the model exhibiting the minimum sum of objectives across the 40 hall of fame models) shows that the error for the entire training set across models is typically less than 2 standard deviations from the experimental values. Features like resting potential, spike amplitude, time-to-first-spike and spike frequency were captured particularly well (Figure 2a; overview of model performance against experiments across Cre-lines: **Figure S1**). To test the optimization workflow against ground truth, we ran cases recovering model parameters as well as electrophysiological response characteristics starting from a fitted input model (**Figure S2**).

**Figure 2:**
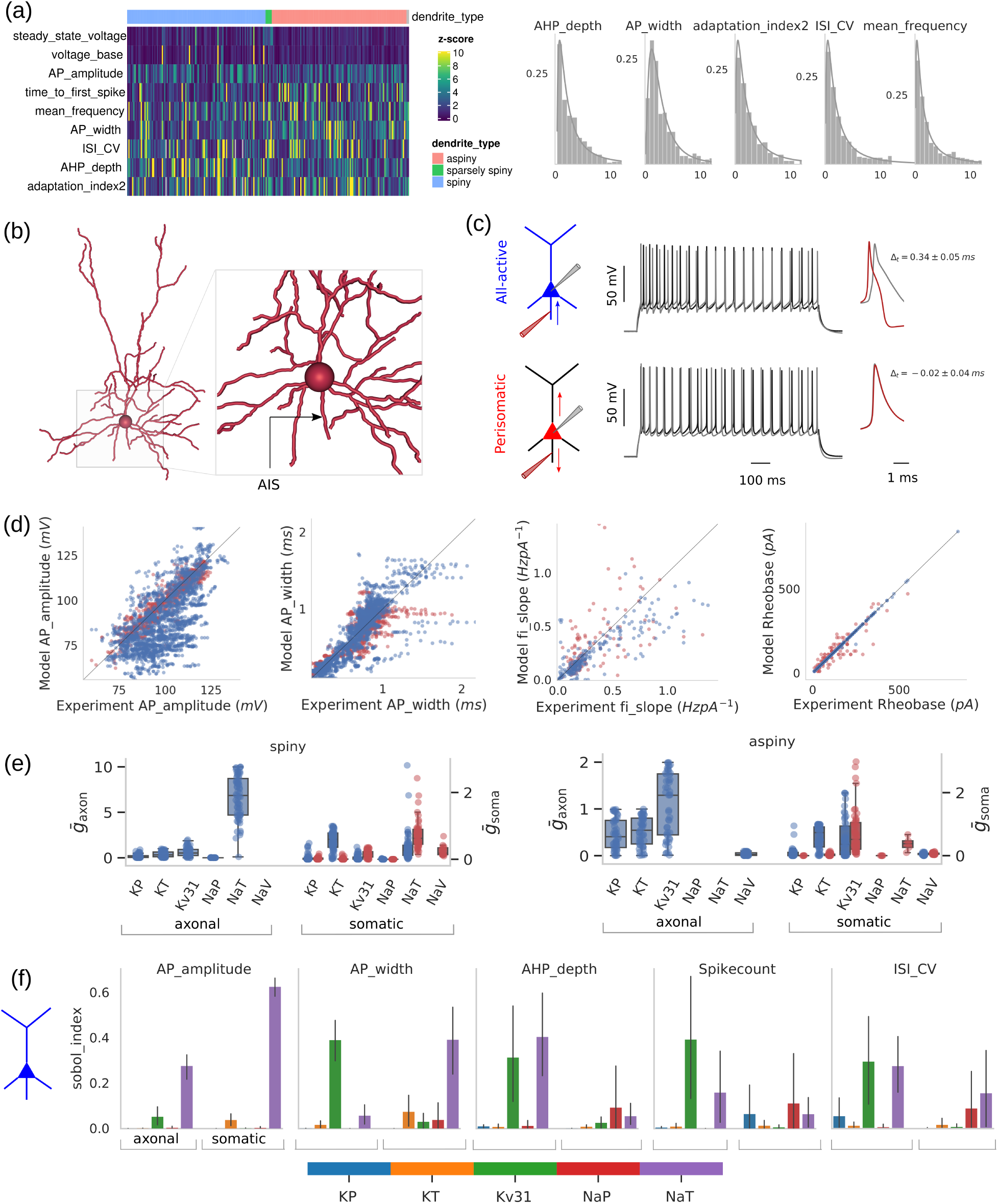
‘All active’ models offer increased bio-realism. **(a)** Performance of 112 spiny, 109 aspiny and 5 sparsely spiny all-active models. (Left) Electrophysiology features for the best models are ranked according to their performance (z-score) during training. AHP depth registered highest training error in most models, whereas voltage-base and steady state voltage are captured accurately. (Right) The histogram of the training error (z-score) for each electrophysiology feature and the associated lognormal fit. **(b)** Reconstructed neuron morphology (CellID: 483101699). **(c)** Both all-active (blue) and perisomatic (red) models capture the experimental (black) somatic voltage response but for all-active models, spikes are initiated from the axon initial segment (AIS) whereas, in perisomatic models, spikes are initiated at the soma. Arrows denote the propagation direction of the action potential. (right) Simulated recordings at axon and soma depict the direction of action potential propagation. 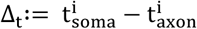, for the i-th spike, >is positive for all-active models and negative for perisomatic models. **(d)** Performance of all models on various electrophysiology features (x-axis: experimentally measured feature; y-axis: model-produced feature after training). Each point corresponds to the feature from a single stimulus protocol. Data along the diagonal of each panel reflect 1-to-1 agreement with experimental values. **(e)** Comparable performance does not translate into similar parameter values. Parameter combinations for all-active models for spiny (left) and aspiny (right) are different from their perisomatic counterpart. **(f)** The relative contribution of AIS and somatic ion channel conductances for each intracellular somatic feature derived from the sensitivity analysis (parameter perturbation: 10% about optimized value; relative contribution of each conductance expressed in terms of Sobol indices). Results shown for all-active models of 10 excitatory cells (bar: mean; error bar: std). While it is known that spike shape features such as spike amplitude and width are critically affected both by AIS and somatic ion channel conductances, AIS conductances are also directly involved with a number of other electrophysiology features such as ISI coefficient-of-variation and spike count.

The all-active models offer enhanced biophysical realism in the form of dendritic and axonal ion channels as well as spikes initiated at the AIS for excitatory cells (Figure 2b-c). To assess the advantages of such enhanced bio-realism, we compared the all-active models with a class of models limiting the presence of ion channels to the soma region, i.e. perisomatic models (^26^; see also **STAR Methods**) that, for example, fail to generate spikes at the AIS. How important is capturing such level of realism? When comparing all-active to perisomatic models originating from the same experiments (thus, representing the same electrophysiology and morphology data) we observe that both model types capture experimental features comparably well (Error! Reference source not found.**a,d**). Yet, preserving the electrophysiological signature using the same set of conductances (including kinetic parameters) by no means implies similarity in the underlying model, i.e. the conductance vectors. In fact, we observe that all-active vs. perisomatic models exhibit substantially different ion channel conductance levels along their morphology. For example, significantly different conductance levels are observed for various Na^+^ and K^+^ ions at the soma for both spiny and aspiny cells (Figure 2e). The same applies for the AIS as perisomatic models, which by design do not account for active conductances anywhere else except the soma.

To what extent do such differences in conductance affect electrophysiology? The fact that both model types exhibit comparable training performance (Figure 2a, d) could suggest that the differences in conductance do not have a measurable effect on electrophysiological features. At the same time, assessing the influence of individual AIS or somatic conductances on a particular electrophysiology (E-) feature is non-trivial as relatively small differences in conductances can significantly affect model response (**Figure S3**). To disentangle the role of individual conductances on specific electrophysiology features, we utilized a method based on polynomial chaos expansions that allows uncertainty quantification and sensitivity analysis to be performed on single-cell models ^36^. The analysis results in a Sobol index for each conductance that represents the contribution of that conductance to the variability of the electrophysiology feature. The Sobol index ranges from 0 to 1 with 0 indicating no contribution of a conductance to a particular E-feature while 1 translates to that conductance explaining all variability with no others contributing. We observe that different conductance vectors between all-active and perisomatic models result in differential contributions of individual conductances to a number of electrophysiology features (Figure 2e).

Specifically, we tested the impact of 5 axonal and 5 somatic ion channel conductances on a set of electrophysiological features using all-active and perisomatic excitatory models from the same experiments (n=10; Figure 2f, **S3**). Despite originating from the same experiments, model predictions on how axonal and somatic conductances affect spike shape and excitability are drastically different between all-active and perisomatic models. For all-active models, the AP-width is equally affected by the axonal Kv3.1 and somatic transient Na^+^ conductance, while perisomatic models predict that somatic Kv3.1 plays the dominant role. In fact, a number of experimental studies have documented the interplay and impact of transient Na^+^ and Kv3.1 channels along the axon during spiking, e.g. ^17,37–39^, offering support to the observations made by the all-active models. Moreover, for AHP depth the all-active models indicate significant contribution both from axonal Kv3.1 and NaT, whereas for perisomatic models the effect remains limited to somatic NaT. Experimental studies have shown that Kv3.1 is expressed in layer 5 mouse pyramidal cells ^40^ and its role in fast AHP ^41^. In general, we observe that the five E-features are affected by a combination of axonal and somatic conductances, whereas the perisomatic models, per design, only account for somatic contributions (often from different conductances). We conclude that the all-active model generation workflow introduced herein, offers a more realistic biophysical mechanism to recapitulate key aspects of experimental electrophysiology compared to perisomatic models.

### 2.2 Model vs. experimental classification of major cortical cell classes

We assessed the quality of the models by the extent to which they captured characteristic properties of a particular neuron or class. Classes separable via a particular electrophysiological, morphological or transcriptomic signature should also be represented by models separable by the same property. To that end, the experimental data we used to generate the single-cell models consisted of a ‘broad’ taxonomy comprising 4 major subclasses (Htr3a, Sst, Pvalb, Pyr). In a further effort to examine the separability of the models at a higher level of resolution (i.e. at a lower level in the hierarchy of cell-type definitions) we used the 19 Cre lines and the 3 dendrite types to define a refined cortical taxonomy. Given the significant overlap of gene co-expression between these lines ^1,2^, we grouped them into ‘refined’ subclasses according to similarity in their gene expression from mouse V1 single-cell RNA-seq data ^1^. This procedure resulted in 11 groups (**Figure S4a**; hierarchical tree dissection at 30%, see **STAR Methods**). From the 11 initial groups, a subset of 6 meaningfully represented transcriptomic cell subclasses (e.g. Oxtr is known to consist of both excitatory and inhibitory cells ^42^ and was thus omitted) had enough experiments across modalities to enable model generation. These 6 subclasses were represented by 4 GABAergic lines namely, Htr3a, Sst, Pvalb, and Ndnf (comprising primarily neuroglia cells) within the inhibitory class, as well as Nr5a1 and Rbp4 representing the L4 and L5 excitatory subclasses within the excitatory class. To assess the utility of the models in preserving cell-type specific identity, we first performed hierarchical clustering (Figure 3a) on the raw feature sets: for the experimental data set we used morphology and electrophysiology (‘Morph+Ephys’) features, whereas for the model data set, we used optimized model parameters (conductance vector) and morphology features (‘Morph+Model parameters’; morphology features same as the ones used in ‘Morph+Ephys’). The close agreement between the two training sets and comparable clustering at the broad subclass level shows that similar information is contained in the experiments and the all-active models generated from these experiments (Figure 3a).

**Figure 3:**
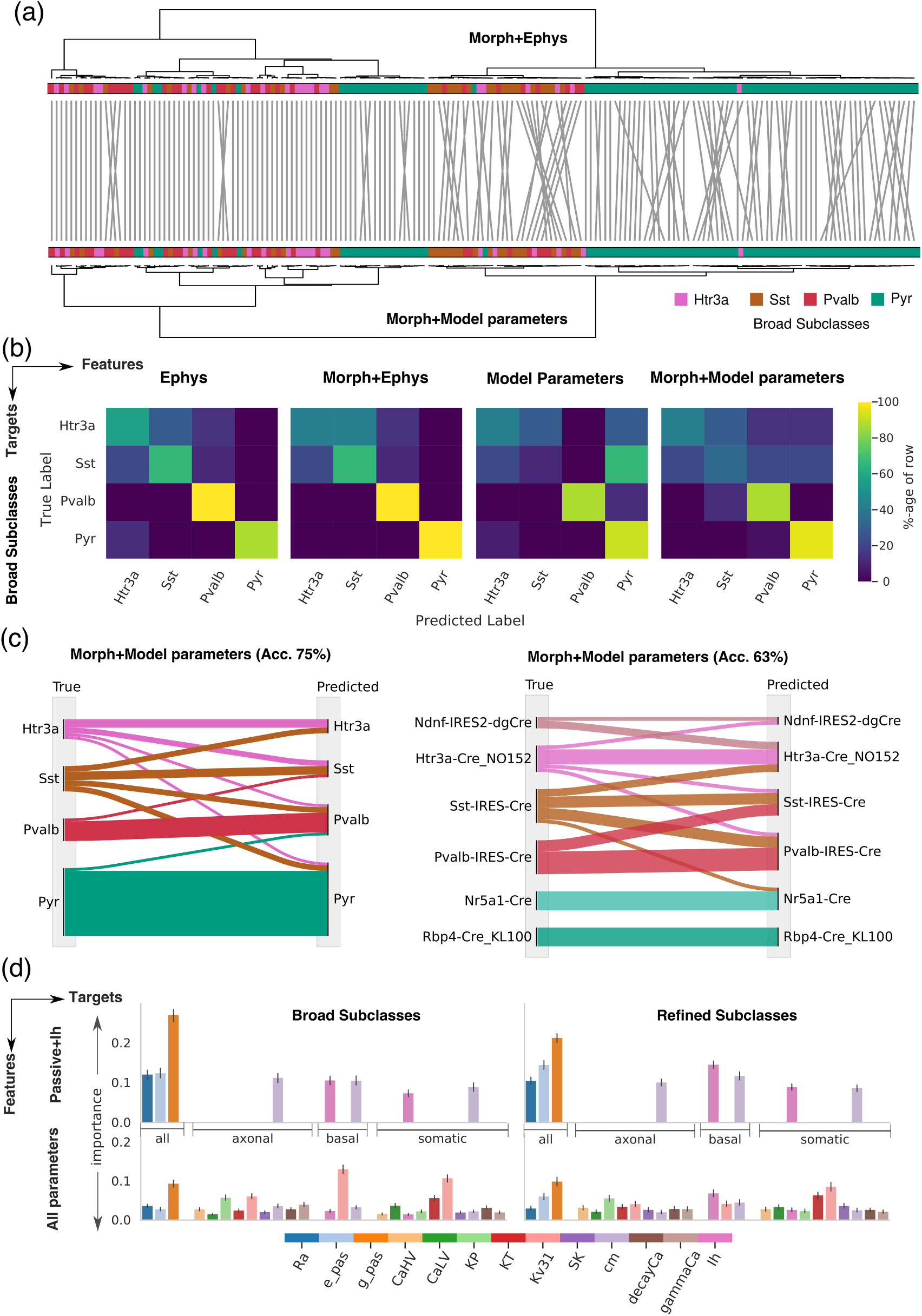
Ion channel conductance specificity along the morphology of all-active models informs of broad and refined subclasses. **(a)** Tanglegram showing the relative agreement between hierarchical clustering based on Morph+Ephys features (top) and Morph+Model Parameter features (bottom) with the colorbar on either side representing the 4 broad subclasses. **(b)** SVM classifier trained on different sets of features (columns) with putative target cell types (rows). The confusion matrix for each feature is shown with test set accuracy 81%, 85%, 69% and 75% respectively. **(c)** Sankey diagrams showing the tendencies for classifiers constructed out of Morph+Model parameters for the two targets on the test set. (left) For broad subclasses Pyr and Pvalb categories are well separated with a few Sst and Htr3a cells misclassified. (right) Clear separation in refined excitatory and inhibitory classes, except for the Sst subclass that is misclassified into an excitatory subclass. **(d)** Random forest classifier trained on model parameters for broad and refined Cre targets as in (b, c). Classifier (top) trained only on passive and h-channel parameters and (bottom) the full set of conductances. Features are sorted according to the parent section of the morphology (’all’-common to the entire morphology, axonal, apical, basal, somatic) and same color is used to indicate a parameter across sections. Among the passive properties (top) *g*_*pas*_ holds the highest importance whereas within the entire parameter set (bottom) Kv3.1 is dominant. Note that going into the full feature set, *p*_*pas*_ retains relative influence in the classifier across targets.

To establish baseline performance metrics, we next assessed the extent to which electrophysiology and morphology features of the experiments can separate between Cre-defined subclasses now in a supervised manner. We trained classifiers (Support Vector Machine with gaussian kernel, see **STAR Methods**) on two feature sets, the experimental electrophysiology (‘ephys’) and the combined electrophysiology and morphology (‘Morph+Ephys’) features. We also trained classifiers on two model-based data sets: first, we trained solely on conductance vectors (‘Model parameters’) and, second, we trained on conductance vectors together with morphology features (‘Morph+Model parameters’; Figure 3b, **S3a**). We observe that while classifiers trained purely on conductance vectors (69%) are less accurate than electrophysiology features (81%), they are both significantly higher than chance (35% and 48%, respectively). In addition, classifiers trained on combined model conductances and morphology features (75%) approach performance with classifiers on experimental electrophysiology and morphology features (85%). Excitatory neurons (Pyr) and the most abundant inhibitory subclass (Pvalb) are clearly separated with high degree of accuracy while the two other inhibitory subclasses, Sst and Htr3a, exhibit wider dispersion (Figure 3c). Classification on the refined taxonomy (6 groups) is also in close agreement as the excitatory subclasses defined in two groups (Nr5a1, Rbp4), are identified with high confidence together with the inhibitory Pvalb (Figure 3c). Inhibitory subclasses such as Ndnf, Sst and Htr3a on the other hand exhibit higher dispersion, contributing to reduced classification performance. We conclude that conductance vectors are indeed informative about transcriptomically-defined cell classes, especially for excitatory and well-separated inhibitory classes such as Pvalb. In addition, we observe that conductance vectors together with morphology features constitute a reliable estimate of cell subclass identity, in keeping with purely experimentally measured features.

Given the discriminative power of these conductance vectors, which parameters or combinations of parameters drive classification? We fitted a classifier on the same cells with broad and refined Cre targets, but now the feature set was constructed solely out of the model parameters (random forest classification). We tested two feature sets: a reduced set comprising of the passive and h-channel conductances (9 parameters), which determine the subthreshold response of the cell and the full shared conductance vector (25 parameters). For the reduced feature set we find that the conductance with the highest classification weight is the passive membrane conductance (g_pas_) with another passive parameter, passive reversal potential (e_pas_), also contributing substantially to classification (Figure 3d). When looking at the full conductance vector, from the active conductances, the two that consistently are weighted high are the basal h-channel conductance and somatic Kv3.1, i.e. two conductances known to contribute in shaping the electrophysiology and response properties of excitatory (h-channel) 43 and inhibitory (Kv3.1) ^39^ cell classes. Encouragingly, the passive and h-channel parameters retain their influence in the full feature set, illustrating their importance in determining cell subclasses.

If passive and h-channel parameters are so important, how necessary is capturing active conductances along the morphology for classifying cells? Notably, the first two stages of our optimization workflow involve optimizing for passive (Stage 0) and h-channel (Stage 1) conductances along the morphology using subthreshold E-features. The first two stages demand a fraction of the computational resources compared to the final optimization stage (Stage 3). We observe that these much reduced conductance vectors offer identical classification performance (**Figure S4c**) at the broad subclass level (69% vs. 69%), but as we traverse deeper in the cell-type taxonomy, represented by the refined classes, the classifier trained on the full conductance set outperforms the one trained on the reduced conductance set (53% vs. 40%, respectively). Notably, the most salient properties of the classification remain unperturbed, e.g., excitatory neurons are better captured than inhibitory neurons while inhibitory Pvalb neurons remain clearly separated. In general, we find that the structure and metrics of the confusion matrices between all-active and passive+h-channel conductance vectors are similar and the governing parameters (g_pas_ and e_pas_) are conserved. We conclude that basic passive parameters as well as I_h_ can distinguish high-level classes whereas the entire spectrum of active conductances better reflects the nuances at finer divisions in the taxonomy.

### 2.3 From electrophysiology to gene expression: model predictions and experimental validation

The all-active models we developed are solely optimized based on electrophysiological features and morphological reconstructions. To what extent can these models reflect the transcriptomic signatures of the cell classes or subclasses they represent? For example, do the conductances derived from the models correlate with expression of specific ion channel genes (notably, the models were not trained on data relating to gene expression)? We looked in the three inhibitory cell subclasses represented by intersection of Cre lines Htr3a, Sst and Pvalb and morphology in the single-cell RNA-seq data set ^1^ (filtered for basket cells in Pvalb, and neuroglia along with bipolar Vip+ cells in Htr3a line; see **STAR Methods**), and focus on the expression profile of K^+^ conductance-related genes, namely, Kcnc1, Kcna1-3,6 Kcnd1-3 associated with Kv3.1, KP-persistent K^+^ and KT-transient K^+^ conductances, respectively, accounted in our models (Figure 4a). Separately, pairwise comparison between the 15 somatic ion channel conductances accounted in the inhibitory all-active models reveals that Kv3.1 conductance is significantly different between models of the three inhibitory subclasses (Figure 4b). The model prediction is that Kv3.1 conductance increases from Htr3a to Sst to Pvalb, with Htr3a exhibiting the lowest and Pvalb the highest Kv3.1 conductance between the three subclasses. When looking at the expression level of Kcnc1, the gene encoding for Kv3.1 ^44–46^, there is agreement between the experimentally measured differential expression and the model prediction between the three inhibitory cell subclasses, i.e. Htr3a < Sst < Pvalb (Figure 4b). Importantly, the initial conditions for all conductances in the optimization of the inhibitory models were identical. We conclude that for the inhibitory cell subclasses considered here, ion channel conductance vectors inferred from the electrophysiology and morphology features are consistent with gene expression patterns for these subclasses, i.e., the conductance vectors also reflect a data modality they did not train on.

**Figure 4:**
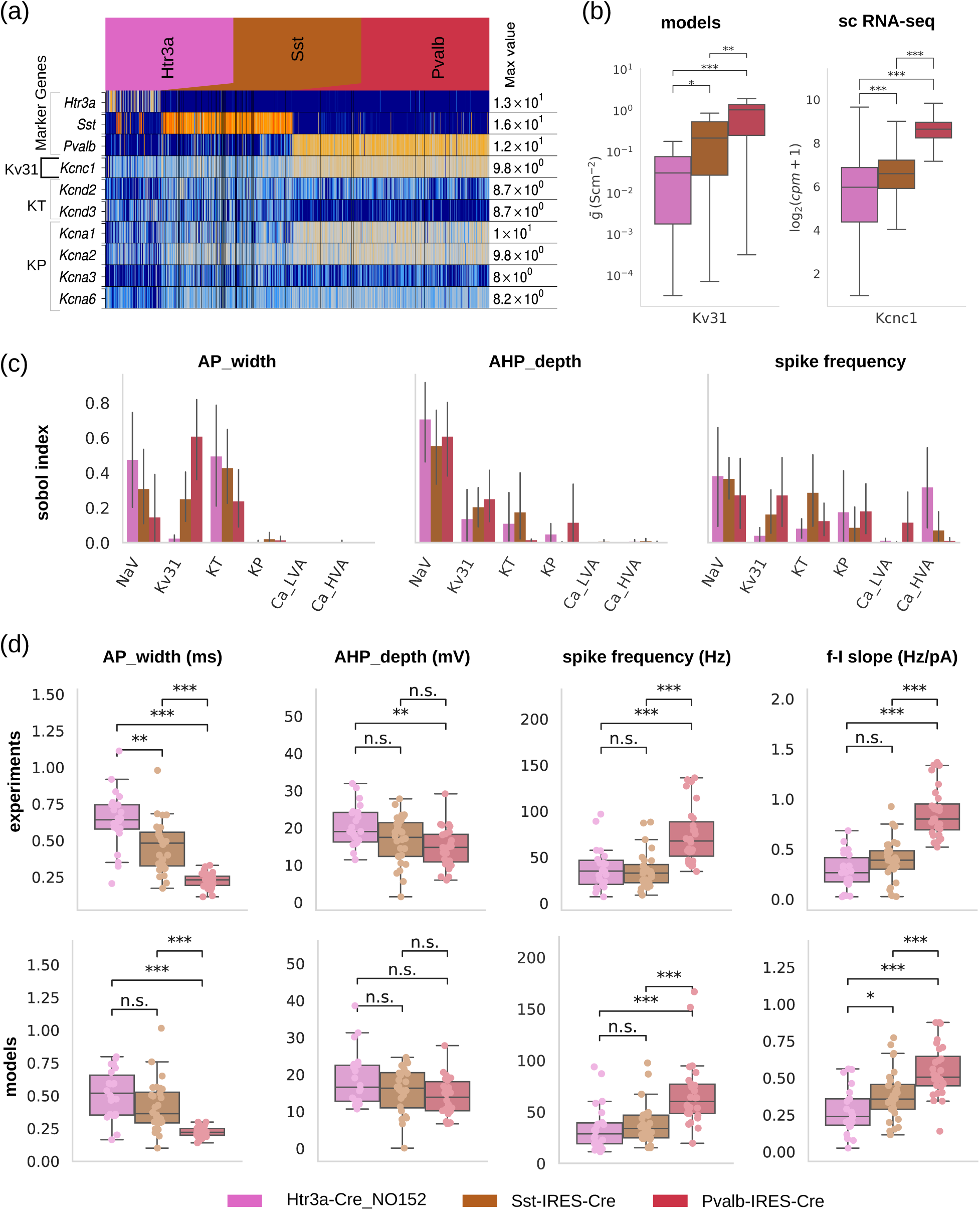
GABAergic cell type differences in voltage-gated ion channel conductances predicted by models are reflected on experimentally measured single-cell gene expression differences. **(a)** Expression profiles of cell type-specific marker genes (Pvalb, Sst, Htr3a) and specific ion channel (Kv3.1, KP - persistent K and KT - transient K) genes (Kcnc1, Kcna1-3,6 Kcnd1-3, respectively) for a set of GABAergic lines in mouse V1 via single cell RNA-seq. **(b)** Comparison between single-cell model, somatic ion channel conductance profiles and single-cell expression of associated genes via single-cell RNA-seq for Kv3.1 (associated gene: Kcnc1); boxplot: median, 1.5 inter-quartile range; cpm: counts per million. Pairwise comparison reveals that somatic ion channel conductance levels of models accurately predicts the direction of type-specific expression profiles (up-vs. downregulation) measured via RNA-seq. **(c)** Ion channels predicted by the models and confirmed by sequencing to differentially express between inhibitory cell types impact intrinsic electrophysiological properties. Sensitivity analysis of single-cell models predicts that beyond somatic Na^+^-conductances, somatic Kv3.1 affects the excitability of neurons (i.e. total number of spikes) as well as spike shape properties such as spike width and AHP depth (line: mean; error bar: 95% confidence interval). **(d:** top) Analysis of electrophysiology properties from *in vitro* experiments exhibits systematic, cell class-specific differences in cellular excitability (f-I slope in response to DC current injection of increasing amplitude) and spike shape characteristics (spike width and AHP depth at 60 pA above the rheobase for each cell). (bottom) Class specific characteristics are preserved in the corresponding all-active models, providing a validation step for the ion channel predictions (boxplot, circles: single cell data). Statistical testing: Mann-Whitney U-test; Statistical significance: *: p-val < 0.05, **: p-val < 10-2, ***: p-val < 10-3, adjusted for a False Discovery Rate (FDR) of 5%.

Can the differences in specific conductances predict the distinct electrophysiological properties of cell types? Given the all-active models of Htr3a, Sst and Pvalb capture relative differences in conductance levels of Kv3.1, we selected three E-features previously shown to have strong relationships to Kv3.1 and tested these relationships with the models. Specifically, we used the Sobol-based sensitivity analysis to assess the impact of 6 somatic conductances (3 of them K^+^-related) on AP width, afterhyperpolarization (AHP) depth and spike frequency (intracellular somatic stimulation waveform: 1s long dc current; Figure 4c). Models predict that Kv3.1 significantly impacts the AP width together with Nav and KT. The AHP depth is mostly affected by the Nav and, more weakly, by the Kv3.1 conductance with Nav mostly influencing Htr3a neurons. Spike frequency is affected by a spectrum of conductances in varying degree. Notably, these results are in line with experimental studies showing the role of Kv3.1 in high-frequency firing ^17,18,39^ and modulation in spike shape (especially narrow spike width and fast AHP ^47–49^) as well as differential expression of Kv channels between Pvalb and Sst subpopulation ^50^. It follows that while multiple conductances affect the chosen E-features, pairwise conductance comparison between models reveals the particular role of Kv3.1 and KT between the three cell types, strongly pointing to causal relationships between transcriptomic and electrophysiology modalities through the latent conductance variable.

Cross-class conductance differences lead to differences in electrophysiological features between classes, i.e. each subclass possesses a specific electrophysiological signature across a set of related features. Experimentally, pairwise comparison of the 4 E-features clearly differentiates the three inhibitory subclasses: Pvalb exibits characteristically tight AP waveform (AP width, AHP depth) and rapid firing (spike frequency, f-I slope) while Htr3a exibits wider AP waveform and slower firing resembling excitatory neurons (Figure 4d, top row). The model simulations performed under identical conditions as patch-clamp experiments (i.e., dc stimulus amplitude and duration; Figure 4d, bottom row) show close resemblance to the experimental data, with absolute values of the E-features as well as between-class relationships conserved between experiments and models. For example, the relationship for the AP width (Δt) is conserved (Δt(Pvalb)<Δt(Sst)<Δt(Htr3a)) between experiments and models with Pvalb exhibiting the smallest AP width (median, inter-quartile range; models: 0.22, 0.05 ms, experiments: 0.23, 0.06 ms), then Sst (models: 0.36, 0.23 ms, experiments: 0.42, 0.16 ms) and, finally, Htr3a (models: 0.52, 0.30 ms, experiments: 0.63, 0.31 ms). Similar comparisons for AHP depth, spike frequency and f-I slope further indicate the close agreement between experiments and models. We conclude that the single-cell models of 3 inhibitory subclasses (i) link electrophysiology and morphology data (model training set) to specific conductance profiles they did not train on, predicting the expression profile of specific ion channel genes as measured by single-cell RNA-seq, (ii) link conductances differentiating between the major inhibitory subclasses with key electrophysiological features that also separate these subclasses, and (iii) capture between-class electrophysiological differences as well as within-class variability as measured in the electrophysiology response of experiments.

We also tested the extent to which similar observations can be made for excitatory cell classes. Specifically, in our refined subclass set we focused on two glutamatergic Cre lines, Rbp4 and Nr5a1, separated based on their transcriptomic profile (**Figure S4a**) with their ion channel-related gene expression ^1^ (Figure 5a,b). Pairwise comparison of conductances between the two excitatory subclasses point to a significant difference in the h-channel conductance with Rbp4 exhibiting larger conductance than Nr5a1 (Nr5a1: n=17; Rbp4: n=17; Figure 5c, left). To test whether the difference predicted by the models is also observed in the expression profile, we examined expression of genes related to the h-channel conductance, i.e., HCN1, HCN2 and HCN3. Pairwise comparison of gene expression for the three genes reveals that for two of them, HCN2 and HCN3, expression in Rbp4 is significantly larger than in Nr5a1 while for the third gene, HCN1, expression is statistically non-significant yet it exhibits a positive effect size in favor of Rbp4 cells (Figure 5c, right).

**Figure 5:**
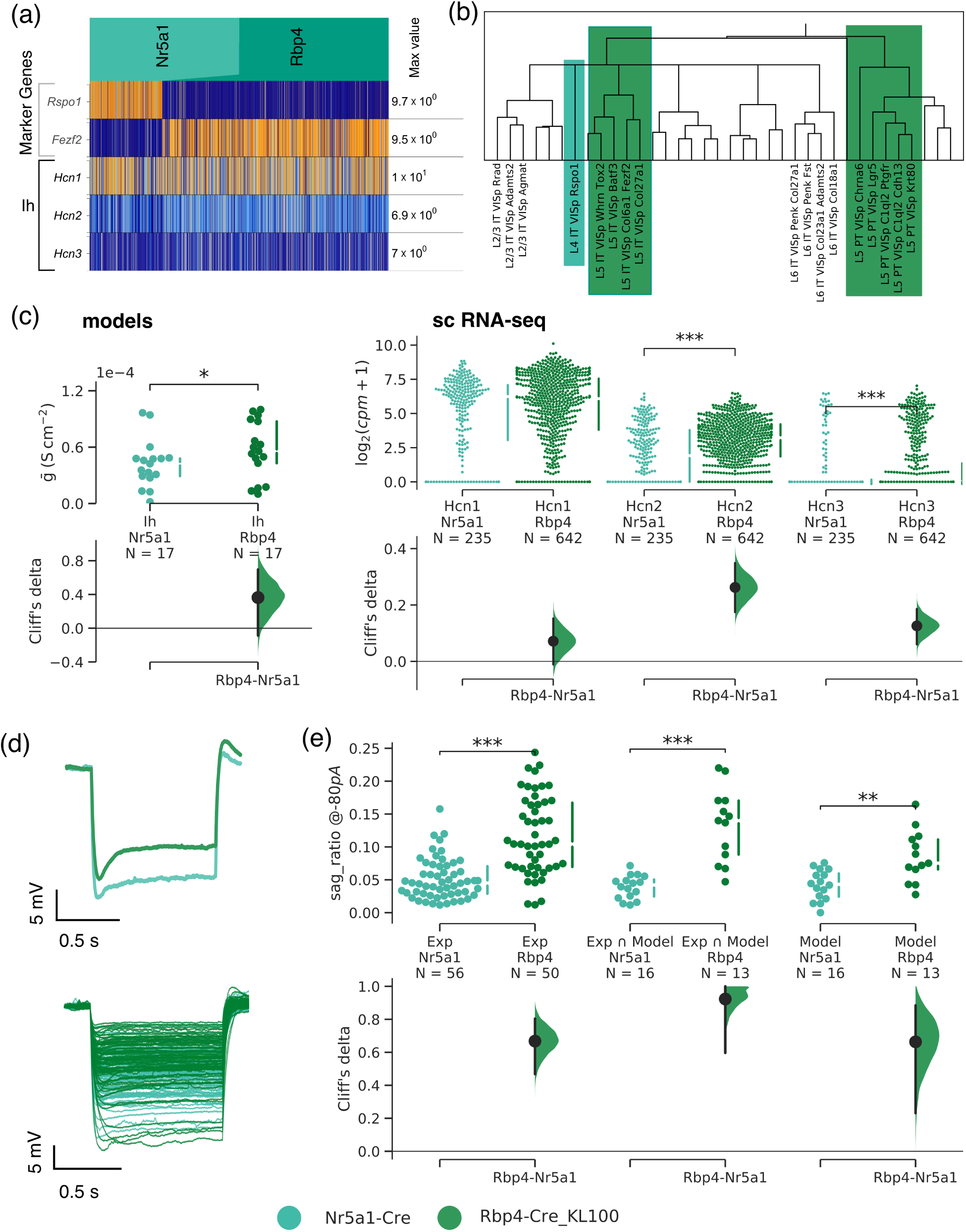
Models predict Ih differences between two excitatory cell classes lead to separation in transcriptomic and electrophysiological space. **(a)** Gene expression for I_h_ channel encoders HCN1-3 along with the marker genes for the two excitatory classes belonging to Cre-lines Nr5a1-Cre, Rbp4-Cre_KL100 (heatmap). **(b)** The two excitatory classes used in the comparison are highlighted in the transcriptomic tree derived from single cell RNA sequencing data ^1^. **(c:** left) h-channel conductance density comparison between Nr5a1 and Rbp4 models shows statistical significance in favor of Rbp4 (p−val<0.05, one-sided Mann-Whitney U-test; Cliff’s delta effect size analysis of the medians ^70^). (right) Same comparison for gene expression from the single cell RNA sequencing data between Nr5a1 and Rbp4 is compatible with model prediction, i.e. Rbp4 exhibits elevated HCN expression compared to Nr5a1. **(d)** Hyperpolarizing traces at −80pA from experiments (top: two cells, bottom: all cells) in the two selected Cre-lines. **(e)** Analysis of sag in the electrophysiology responses. Sag ratio (**Table S2**) comparison between (left) all cells of the two Cre-lines, (middle) the subset of experiments for which models were generated, and (right) the corresponding model responses. The trend revealed by the modeling prediction, i.e. higher h-channel conductance density in the Rbp4 vs. Nr5a1 line, holds across comparisons.

We then asked whether the difference in the h-channel conductance between Rbp4 and Nr5a1 neurons can explain differences in their electrophysiology. The feature robustly linked to h-channel is the sag response to a hyperpolarizing current injection ^51^ (Figure 5d). When looking at the response quantified by sag_ratio (**Table S2**) to such hyperpolarizing injections of Rbp4 vs. Nr5a1 cells, Rbp4 cells exhibit significantly larger sag-response than Nr5a1 (Figure 5e left, middle) for all experiments as well as the subset of the modeled cells. When looking at the models, we also see a sag-response significantly more pronounced for Rbp4 than for Nr5a1 models, an outcome of their increased h-channel conductance. We summarize that the main conclusions drawn from comparisons between inhibitory subclasses also hold for models of excitatory subclasses represented here by Rbp4 and Nr5a1-Cre lines.

### 2.4 Model uniqueness: within vs. across cell class variability

So far, a major assumption of our approach has been that a single model adequately captures features of a single-cell experiment. This ‘best model’ is the outcome of the optimization workflow and we use it to classify cells with respect to characteristic properties and link across data-modalities. On the other hand, generating conductance-based single-cell models from electrophysiology data is an ill-posed optimization problem that can result in widely different solutions for a particular experiment ^7^. How much does that affect the models generated herein and their interpretation? Addressing this question is particularly important given that the experimental data and associated models span across cortical layers, morphology and electrophysiology types, i.e. exhibit considerable within-as well as across-cell type variability.

To address this point we looked beyond the ‘best model’ of the optimization and, instead, study the 40 best models exhibiting the smallest sum of objectives (“hall of fame” or hof models) out of the optimization workflow ^34^. These hof models are representations of the E-features of a single-cell experiment given a reconstructed morphology (Figure 6a), constituting the group of best models out of more than 600,000 models evaluated during the optimization. How much better does the ‘best model’ perform over its hof counterparts? A substantial decrease in model performance would point to the best model representing a unique and near-optimal solution in the parameter landscape. Instead, when looking at the performance of the hof models we observe a graded increase in the training set error (z-score). Moreover, hof models perform equally well (measured in terms of explained variance, see **STAR Methods**) for noisy stimulation they were not trained on (Figure 6b). In other words, while the best model of an experiment represents a solution with the smallest training error, the 39 other hof models generalize equally well in two different types of stimulation sets irrespective of their initialization (**Figure S5b**). This points to a single experiment reflected in a family of models with similar performance metrics rather than the existence of a single, optimal one.

**Figure 6:**
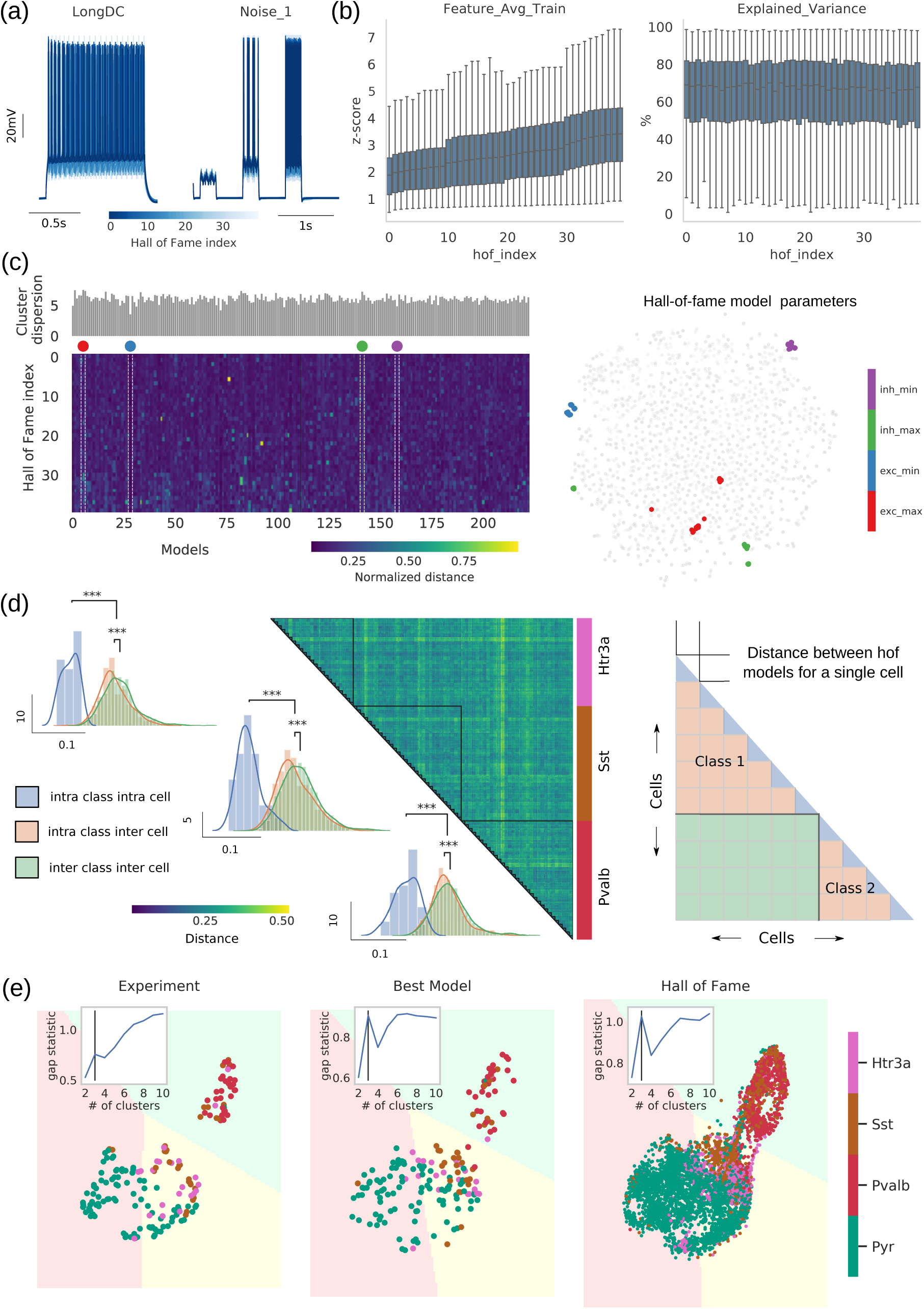
Model representation of broad cell subclasses remains robust in face of biological variability and optimization complexity. **(a)** 40 hall of fame (hof) models are generated for each cell (response of hof models to 1s dc and noise stimulus). **(b)** Feature average (computed in terms of z-score) and explained variance shown for 230 cells against the hof index (sorted according to training error of each cell). While the feature average exhibits an increasing trend (increasing error with hof index), explained variance (to untrained stimuli) is comparable across hof indices. **(c:** left) The hof parameter set for each cell creates a cluster in ñ-dimensional euclidean space, where ñ denotes the common parameters across all cells. The centroid of this cluster is calculated (color: euclidean distance between hof index and its centroid). The sum of all distances is a measure of cluster dispersion (barplot shown on top of the distance matrix). (Right) t-SNE plot highlights four cases of maximally (red: excitatory cell; green: inhibitory cell) and minimally (blue: excitatory cell; magenta: inhibitory cell) dispersed parameter vectors in the reduced 2D space. **(d:** left) Heatmap showing euclidean distances between hof models for the 3 broad inhibitory subclasses - each blockdiagonal represents 40 hof models for a single cell. The schematic (right) illustrates the interpretation for different region of the heatmap, namely, intra cell intra class (blue), inter cell intra class (orange) and inter cell inter class (green). The darker blockdiagonals as well as the distribution of the intra-inter distances for the 3 broad subclasses indicate an ordered structure in the parameter dispersion, i.e., degenerate parameters for a single cell are tightly clustered compared to parameters of the same broad subclass, followed by model parameters between diffferent classes (p-val<0.01; Mann-Whitney U-testing). **(e)** For cells belonging to the 4 broad subclasses we calculate the hof model electrophysiology features at the maximal amplitude stimulus protocol and project them onto UMAP embedding of the corresponding features at the experiment level (optimal number of clusters: *n*_*clusters*_=3 via gap statistic ^69^). The corresponding k-means decision boundary is drawn on the embedded space for the experiment, the best model (hof index=0) and all hof models (40 models per cell). The number of detected clusters, their composition and associated decision boundaries remain unaltered.

Does degeneracy of the solutions i.e., existence of multiple conductance vectors for a single-cell impede our ability to draw insights from models? If wildly different conductance vectors can result to similar E-features and metrics, then our previous observations about such combinations being characteristic of cell class would be put under question. To address this point, we look closer into hof conductance dispersion. Specifically, we measured dispersion of conductance vectors within experiment (hof) vs. across a cell class: if conductance variability of hof models is larger than conductance variability across models of the same cell class, it would be difficult to ascribe association between conductance vectors and cell type identity. We observe that despite variability (Figure 6a, b), conductance vectors in hof models for a particular experiment exhibit significantly smaller dispersion than across any of the four major cell classes (Figure 6c,d, **S5**). To validate that these global cross-class relationships in the models are also consistent in the space of E-features, we compare the clustering structure in this space between experiments, models, and hof population. The clusters detected using experimental E-features (Figure 6e left) are conserved when using the same features from the best model (Figure 6e middle). When features from the hof models are accounted for (40 models per experiment; in total: 230×40=9200 models evaluated), the number of clusters and, more importantly, decision boundaries are in close agreement between experiments, best model and hof models (Figure 6e right). This is consistent with the previous analysis (Figure 3b), which showed that the ephys structure captured by supervised classifiers is preserved within model-based classifiers (the three unsupervised clusters capturing the most salient cross-cluster relationships without class label information provide additional validation). We conclude that conductance vectors generated by the all-active optimization workflow reflect distinct properties of cell classes rather than parameter or experimental variability and conserve the relationship between major cortical classes.

## 3. Discussion

We present a computational optimization workflow for *en masse* generation and evaluation of bio-realistic and morphologically accurate single-cell models with active conductances along their entire morphology (‘all-active’) for more than 230 *in vitro* experiments from mouse visual cortex. We use this workflow to evaluate 130 million models across experiments and select either the best model or a population of best models (“hall-of-fame”) to represent measurements from single-cells. The number of models produced allowed us to integrate across cell-type taxonomies and was key to establishing causal links between modalities: morphology, electrophysiology and transcriptomics. We account for 15 passive, h-channel, Na^+^, K^+^ and Ca^++^ conductances known to exist in cortical neurons and use the optimization to fit their value along the entire morphology. Evaluation of the models exhibits their ability to capture important electrophysiological features of cortical neurons such as axonal spike initiation and points to the existence of distinct ion channel conductance vectors (i.e. conductance values along the neural morphology) that distinguish between major cortical classes.

The all-active models indicate that the passive membrane conductance g_pas_ and reversal potential e_pas_ as well as h-channel conductances are of particular importance in classifying major cortical classes. We show that at two different levels of cell subclass (broad and refined), these two conductances are important in determining the membership of a particular cell within a subclass. In terms of coarse classification (Htr3a, Sst, Pvalb and Pyr), using only the passive and h-channel conductance combination is as informative as using the entire conductance vector. This means that, given the herein introduced model generation workflow, only the first two stages of the workflow are required to generate models with properties accounting for the broad subclasses, i.e. the computational demand is reduced by orders of magnitude (**Figure S6-7**). In terms of classification, this translates into the majority of broad subclasses being well-separated so that basic conductance and morphology properties suffice to distinguish between them. (Notably, morphology properties are bound to affect the conductance vector in the optimization.) As we traverse deeper into the cell-type hierarchy going from a broad to refined subclasses (4 to 6), the random forest classification performance increases in favor of the full conductance vector (**Figure S4**). In other words, it is when trying to distinguish between finer cell classes that active conductances impact classification and the importance of conductance combinations details becomes evident. It follows that the performance of the classifier is currently limited by the training data size (230 conductance vectors) – just as electrophysiology features are shaped by a spectrum of conductances rather than the value of a single one, we predict that with a larger training set a model-based classifier could leverage such combinations for performance well above the baseline set by the passive, h-channel combination.

We show that all-active models preserve between-subclass relationships despite biological and experimental variability and offer two lines of support: (i) classification on a feature set constructed on conductance vectors and morphology properties, and (ii) within a low dimensional embedding of the electrophysiological feature space, the decision boundaries are preserved between broad subclasses for experiments and models (including hall of fame models). We expect that Patch-seq 52,53, where one can obtain morphology, electrophysiology, and transcriptomics for the same cell (as opposed to relying solely on Cre-reporter lines), will lead to more accurate mapping between model parameters and cell-types in transcriptomic space.

While pharmacology and genetic manipulations offer unparalleled insights on the link between genotype and phenotype, this process remains laborious, expensive and time-consuming. On the other hand, an important aspect of models is their ability to link seemingly disparate observations – in modeling, all parameters are accessible to the experimentalist. Can the all-active models inform about the gene profile of cell classes they are supposed to capture? More typical model generation approaches that focus on one or a small number of cell types are limited in their ability to answer such general questions. By generating a large population of all-active models, we show that for a set of excitatory and inhibitory classes, differences between models solely trained on electrophysiology predict differentially expressed genes related to specific conductances. These results are experimentally validated by single-cell RNA-sequencing. Specifically, between three major inhibitory classes, Pvalb, Sst and Htr3a1, all-active models predicted differences in Kv3.1 (ḡ_Pvalb_>ḡ_Sst_>ḡ_Htr3a1_) validated by transcriptomics. Additionally, sensitivity analysis shows that these conductances impact key electrophysiology features such as the AP width, AHP depth, spike frequency and f-I slope. Moreover, we show that these electrophysiology features, indeed, separate the three inhibitory classes offering a causal relationship between the differences observed at the genetic level (i.e., genes related to Kv3.1) and the functional level (i.e., cell classes separated by sets of electrophysiology features). The same approach yielded differences between two excitatory subclasses, Nr5a1 and Rbp4, revealing the connection between the h-channel conductance (ḡ_Rbp4_ > ḡ_Nr5a1_) and expression of HCN genes.

To establish links between data modalities we have used the channel conductances of models generated by the optimization to identify gene expression differences at a cell class level. Such validation is not always straightforward, especially when a group of genes encodes subunits of a single channel, e.g., for the inhibitory classes reconciling the KT (transient K^+^) conductance densities and the expression of the responsible genes remains a future challenge. While our models assume the existence of 15 ion channel conductances shown to be important for cortical neurons, it is also clear that unless an ion channel conductance is included in the initial recipe fed into the optimization framework, it will not be accounted for in the analysis. On the other hand, inclusion of an increasing number of conductances at the model-generation stage to account for the entirety of relevant channels (e.g. given recent model/data availability ^54^) renders the optimization increasingly challenging despite recent advances ^55–57^. A solution to this challenge may include an initial step where the conductance recipe gets informed by the class-specific profile from single-cell sequencing (e.g., RNA sequencing, ^1,2^) which, in a second step, is fed into the optimization framework to define the conductance values. Such an approach would result in class-specific recipes that would be refined by single-cell models allowing for more elaborate sensitivity analysis and parameter exploration. Notably, in our model generation routine this will overrule a key assumption – uniform conductance bounds across experiments at the algorithm initiation, which was imperative to our interpretation of emergent separation at the ion channel level between cell classes.

In our effort to establish the relevance of a low-dimensional representation of the experimental data modalities vis-a-vis conductance combinations capturing cell-type specific information and translating these differences into different modalities (transcriptome and electrophysiology), we analyzed the uniqueness of the derived models. The degeneracy of biological systems is well established ^58,59^. In a highly constrained multi-objective optimization formulation, we evaluate the dispersion between models from the same experiment. For the best performing models across different seeding of the initial model population we found that intra-cell separation (dispersion between conductance vectors for the same cell) is significantly lower than the inter-cell separation (dispersion of conductance vector pairs from different cells) within the same broad cell subclass. Furthermore, the separation between models at inter-class level is significantly higher than both intra class metrics. The outcome of this is that models of broad cell subclasses preserve their cell-specific identity. We postulate that this result is a direct consequence of the constraints introduced in the optimization, which is supported in recent finding within a neural density estimator framework 57. We conclude that all-active models predict/capture molecular differences offering a systematic and causal way of linking evidence across data modalities and cell classes.

## 4 STAR Methods

### 4.1 Slice electrophysiology and morphology reconstructions

The electrophysiology and morphology data are part of the Allen Cell-Types Database 60. Whole cell patch clamp recordings were performed on single neurons in prepared slices from adult mouse visual cortex. Fluorescence illumination was used to identify the Cre-positive neurons based on tdTomato fluorescence. Time series for the voltage response under stimuli of varied amplitude and duration for standardized set of protocols (step, ramp, colored noise) are collected in an. nwb 61 format at 34∘C and corrected for liquid junction potential (−14mV). Following the recording protocols the neurons are filled with Biocytin, present in the electrode. Sections are processed using 3,3’-diaminobenzidine (DAB) peroxidase substrate kit to identify recorded neurons filled with biocytin. Serial images (63X magnification) through biocytin-filled neurons were evaluated for quality, and cells that passed a quality threshold are entered a detailed morphological analysis workflow. The dendritic morphology of each neuron is identified as either aspiny, sparsely spiny or spiny. Reconstructions are generated for a subset of neurons using a 3D Visualization-Assisted Analysis (Vaa3D) ^29^ workflow. The automated 3D reconstruction results were then manually curated using the Mozak extension of Vaa3D and subsequently saved in the swc format. The electrophysiology and morphology procedures along with the immunohistological staining are detailed in the technical white papers of the cell-types database ^60^.

### 4.2 Single cell RNA sequencing and expression analysis

Materials and methods used for single cell RNA collection and sequencing are described in ^1^. Briefly, single cells from visual cortex of adult transgenic mice were labeled and isolated by FACS (fluorescence activated sorting). SMART-Seq Ultra Low Input RNA Kit for Illumina Sequencing was used for cDNA synthesis of single-cell RNA and subsequent amplification. Further downstream analysis involved aligning reads and QC. This data, consisting of more than 1600 cells, is freely available through the Allen Brain Atlas data portal (http://www.brain-map.org).

The broad subclass definition used throughout this paper consists of 3 primary inhibitory lines – Pvalb basket cells, Sst, Htr3a – neuroglia and bipolar Vip cells, with the fourth excitatory class ‘Pyr’ is composed of spiny Rorb-Ires2, Scnn1a-Tg2,3, Nr5a1 and Rbp4 cells. To infer the refined classes, we performed hierarchical clustering of the gene expression correlation matrix, which is computed by mapping the t-type to t-type correlation to Cre-Cre correlation and averaging them over multiple trials in a Monte-carlo fashion. The clusters were selected by dissecting the hierarchical tree at 30% of the maximum distance resulting in 11 groups namely, Vipr2-IRES2-Cre, Ndnf-IRES2-dgCre, Nos1-CreERT2, (Pvalb-IRES-Cre/Nkx2-1-CreERT2), (Sst-IRES-Cre/Chrna2-Cre_OE25), Oxtr-T2A-Cre, (Htr3a-Cre_NO152/Vip-IRES-Cre), Ctgf-T2A-dgCre, (Slc17a6-IRES-Cre/Tlx3-Cre_PL56/Scnn1a-Tg2,Tg3-Cre/Nr5a1-Cre), Esr2-IRES2-Cre, (Rbp4-Cre_KL100/Ntsr1-Cre_GN220). For classification and comparison between classes, a target subclass, broad or refined, needed to have at least 10 member cells. To ensure maximal discriminability we also remove Cre lines with overlapping expression patterns so that the initial 11 clusters (**Figure S4a**) were reduced to 6 refined classes – Ndnf, the 3 inhibitory lines form the broad subclasses (Pvalb, Sst, Htr3a), and Nr5a1, Rbp4 from the excitatory group. Note that these definitions are used for other downstream analysis throughout this paper – comparison of 3 primary inhibitory lines (Figure 4) and comparison of two excitatory lines (Nr5a1, Rbp4) from refined class definitions (Figure 5).

In reconciling the two separate data sets, the transcriptomics, single-cell RNA-seq set ^1,2^ with the single-cell morphology and electrophysiology (ME) set ^8^, we performed additional filtering on top of the Cre-lines to ensure consistency in comparing classes across the two datasets. First, we labeled Cre-lines in the ME data which are not sampled in the RNA-seq data as ‘Other’ (Figure 1a, b) and not used them in any cell class definitions. The only exception being Rorb-Ires2-Cre which is part of the Pyr class, since a significant portion of these cells were sampled during model generation. For the analysis of the inhibitory classes, we sought to keep the comparisons limited to basket cells within the Pvalb class (by filtering out Chandeliers, Pvalb-Vipr2 labeled cells on the T side and qualitative morphology label ‘Chandelier’ on the ME side) and within Htr3a line by keeping neurogliaform (marker gene: Lamp5) and bipolar Vip cells. Similarly, on the excitatory side Rbp4 cells are found across glutamatergic and GABAergic divide in the transcriptomics data. Thus, within the two selected Cre-lines, Nr5a1 and Rbp4, we kept the comparisons limited to L4 intratelencephalic (IT) and a combination of L5 pyramidal tract (PT) and IT cells (Figure 5b) respectively. On the ME side we have simply used the layer assignment (layer 4 spiny Nr5a1 vs layer 5 spiny Rbp4 cells) as the filtering criteria for conductance and E-feature comparisons.

### 4.3 Single cell model generation and analysis

The models are simulated in NEURON 7.5 ^62^ simulation environment via python interface ^63^ with variable time step integration. Due to the inconsistencies in the axonal reconstructions, we have uniformly replaced the axons (irrespective of their presence in the .swc file ^64^) with a 60μm stub with a diameter of 1μm after loading them using Import3d_SWC_read() in NEURON. For model generation we have used a combination of the following active ion channels: I_h_, NaT, NaP, KT, KP, Kv2, Kv3.1, SK, I_m_, Ca_LVA_, Ca_HVA_ for apical, basal dendrites, soma and AIS. At exploratory stages of workflow design, we found that optimizing the entire parameter vector (43 variables) in a single stage led to poorer convergence. We attribute this to the passive properties since models are particularly sensitive to these parameters. To address this point, we introduced a preliminary stage (Stage 0) with only cm, Ra, g_pas_, e_pas_, to optimize the baseline response properties of the model and allow them to be optimized during the successive stages. Next, we observed that lumping h-channel conductance as well as other active channels in a single stage and adding the deviation between the model and experimental sag to the objective vector, the suprathreshold component of the error vector dominated the resultant model generations. This led to the decision to introduce a separate stage for the h-channel (Stage 1) followed by a third stage that includes the rest of the active channels (Stage 2).

We used a 1s long dc current step for eliciting the neural response in experiments and models. In total, we use 15 electrophysiology features over 3 stages: resting potential before and after current injection (voltage base, steady state voltage), average membrane potential during stimulation (voltage deflection), spike frequency, ISI slope, adaptation index, time-to-first-spike, AP amplitude and width, AHP depth ^65^. Models are evaluated against the experimental features with respect to 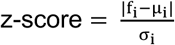, where the feature value f_i_ is measured from the output traces of the models while µ_i_ and σ_i_ are the experimentally measured mean and standard deviation, respectively, for the targeted electrophysiology features. From the definition, when z-score=1, model and experiments are within one standard deviation of the experiment. The experimental standard deviation for each feature is calculated over repetitions of the same stimulus; in absence of repetition we use 5% of the mean as the default standard deviation. During training we use the force_max_score=True option in the efeature module of BluePyOpt such that maximum error, i.e., z-score for any objective between model and experiment is bounded by 250. A common example when this option will be triggered is when in comparison to spiking trace in the experiment the model for the same stimulus produces sub-threshold response.

1. Stage 0: fit passive membrane properties of the model along the entire single-cell morphology on subthreshold, non-spiking features of an experiment for a number of stimulation amplitudes (# of training objectives = 4 × # subthreshold traces; # generations= 50; # seeds = 1),
2. Stage 1: include the h-channel conductance and re-fit along the entire single-cell morphology on subthreshold experimental features (# of training objectives = 5 × # subthreshold traces; # generations= 50; # seeds = 1). We have selected h-channels to include in the intermediate stage to fully characterize the subthreshold responses before exposing the models to the battery of stimulus driving spiking behavior.
3. Stage 2: fit all active Na^+^, K^+^ and Ca^++^ -conductances on subthreshold and spiking features for a number of experimental stimulus amplitudes (# training objectives = 11 × 2 spiking traces + spikecount at rheobase trace and the maximal subthreshold trace; # generations= 200; # seeds = 4).

The list of parameters, their description as well as their bounds and the stimulus set used at each stage to inform the optimization is detailed in **Table S1-3**. Using this model optimization workflow, approximately 600,000 models are tested during the development stage. It follows that this procedure is associated with a significant computational demand, i.e. the CPU demand per cell at the in-house computing cluster is 26 ± 11 hours on a 256 × 2.2 GHz Intel Xeon E5-2630v4 processor rack for a 2:1 offspring to core ratio (we have also leveraged 3 other computing systems, Amazon Web Services EC2, NERSC and Blue Brain 5 Supercomputers, detailed in **Figure S6**). However, we were able to drastically improve the computation demand by adding a timeout functionality to the evolutionary algorithm subroutine, where models for which evaluation surpassed the set timeout parameter are rejected with maximum score across all objectives (**Figure S7**).

The model generation has a final validation stage - where the models go through the entire stimulus battery of the corresponding experiment. In particular we calculate two more metrics along with the average training error (z-score) across all the objectives, namely, Feature average Train: the Feature Average Generalization, which is the average z-score of the 15 features used in the optimization for all the 1s long DC stimulus that were not part of the training set in Stage 2, and finally, the ratio of the trial-to-trial explained variance (EV) ^26,66^ of the spike times for colored noise - a stimulus type the model has not seen at any level of the model generation. This is defined as

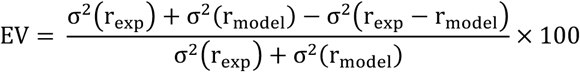

where σ^2^ is the variance, the Peri Stimulus Histogram (PSTH) denoted by r, of the experiment and model are calculated by convolving the respective spike trains with a gaussian curve at a bin window of 10 ms. We report these metrics for all 40 hall of fame models across the 4 seeds.

### 4.4 Classification and Dimensionality Reduction

We have used the built-in Support Vector Machine (SVC(kernel=’rbf’)) in a pipeline on top of StandardScaler() and Random-Forest classifier (RandomForestClassifier()) within Scikit-learn ^67^ library for our classification results in Figure 3. To construct the different combinations of feature vectors we employed the ephys features used in training for model generation, morphological features, and, finally, the model parameter set with least training error. For morphological features we have used the python package NeuroM ^68^ to extract the following features from the input. swc reconstructions - soma_surface, soma_radius, and length, area, volume, taper_rate for all the neurites (tree of sections), i.e., apical, basal dendrite, axon. For model-based feature construction we only consider parameters that are shared by all cells to prevent obvious differences between excitatory and inhibitory cells (e.g., no apical dendrites for aspiny/sparsely spiny cell reconstructions, as well as differences in conductance recipe (**Table S1**) built into the model generation pipeline) from dominating the results. The four feature sets in Figure 3b comprises of 7, 23, 25, 41 features each. ‘Acc’: classification accuracy is reported on a stratified test set with 70-30 train-test split of the data and SVM, random Forest hyperparameters tuned in a 3-fold cross-validation of the training set; arrows: ‘Acc’ reported as increase/decrease over ‘chance classifier’ which are randomly sampled labels from the test set. The confusion matrices for random forest classifier are shown in **Figure S4c**, with the 2 × 2 parameter ‘importance’ grid (averaged over all trees in the ensemble) in Figure 3d. The importance of a feature simply corresponds to average reduction in gini impurity for nodes that uses that specific feature.

In Figure 6e we have used the UMAP transform (github.com/lmcinnes/umap) with n_neighbors_ = 10, again in a pipeline with the scikit-learn StandardScaler(). To enable comparisons between experiments and models for embedded electrophysiological features (at the maximal 1s long DC step current used in training), we learn the manifold on the experimental features and use the transform() method in UMAP to force ‘new data’ (model ephys features) in the same 2D space. We use scikit-learn KMeans() for an unsupervised clustering of the reduced ephys features and consequently draw the decision boundary by assigning cluster tag by sampling uniformly from the 2D grid. The optimal no. of clusters in KMeans is determined using gap statistic ^69^.

### 4.5 Code availability

The automated staged optimization workflow is available as a python package (<u>github.com/AllenInstitute/All-active-Workflow</u>) written on top of BluePyOpt. The workflow is easily configurable using nested .json schemas. The electrophysiology, reconstructed morphology as well as the perisomatic models can be downloaded using <u>AllenSDK</u> api and as a result is a dependency of our repository along with NEURON for biophysically detailed simulations. As a placeholder until a future data release where these new models are integrated into the Allen Institute Cell-types data portal (<u>portal.brain-map.org/explore/models</u>), we have shared the models used in this work in a separate github repository (<u>github.com/AllenInstitute/All-active-Manuscript</u>). The models are consistent with the previously established Allen Institute biophysical format.

## Acknowledgements

We wish to thank the Allen Institute founder, Paul G. Allen, for his vision, encouragement, and support. We also thank Brandon Blanchard for his help in visualizing the mouse brain atlas and Fahimeh Baftizadeh for her input in analyzing single-cell RNA-seq data. The BlueBrain 5 and Linux cluster used as a development system for this work is financed by ETH Board Funding to the Blue Brain Project as a National Research Infrastructure and hosted at the Swiss National Supercomputing Center (CSCS). This research used resources of the National Energy Research Scientific Computing Center; a DOE Office of Science User Facility supported by the Office of Science of the U.S. Department of Energy Office of Science User Facility operated under Contract No. DE-AC02-05CH11231.

## Notes

https://github.com/AllenInstitute/All-active-Workflow

### Competing Interest Statement

The authors have declared no competing interest.

