## Supplemental material for "Single-neuron models linking electrophysiology, morphology and transcriptomics across cortical cell types"

### Supplementary Material

| Parameters | Apical Dendrite | Basal Dendrite | Soma | Axon | Unit | Modfile |
|---|---|---|---|---|---|---|
| $c_m$ | $[0.5, 10]$ | $[0.5, 10]$ | $[0.5, 10]$ | $[0.5, 10]$ | $\mu Fcm^{-2}$ | pas |
| $g_{pas}$ | $[10^{-7}, 10^{-2}]$ | $[10^{-7}, 10^{-2}]$ | $[10^{-7}, 10^{-2}]$ | $[10^{-7}, 10^{-2}]$ | $Scm^{-2}$ | pas |
| $e_{pas}$ | $[-120, -60]$ | $[-120, -60]$ | $[-120, -60]$ | $[-120, -60]$ | $mV$ | pas |
| $R_a$ | $[50, 200]$ | $[50, 200]$ | $[50, 200]$ | $[50, 200]$ | $\Omega cm$ | pas |
| $I_h$ | $[10^{-7}, 10^{-4}]$ | $[10^{-7}, 10^{-4}]$ | $[10^{-7}, 10^{-4}]$ | - | $Scm^{-2}$ | Ih.mod |
| NaT | $[0, 10^{-1}]$ | $[0, 10^{-2}]$ | $[0, 5]$ | - | $Scm^{-2}$ | NaTs2_t.mod |
| NaT | - | - | - | $[0, 10]$ | $Scm^{-2}$ | NaTa_t.mod |
| NaP | - | - | $[0, 1]$ | $[0, 5]$ | $Scm^{-2}$ | Nap_Et2.mod |
| NaV | - | $[10^{-7}, 10^{-1}]$ | $[10^{-7}, 10^{-1}]$ | $[10^{-7}, 5]$ | $Scm^{-2}$ | NaV.mod |
| KT | - | - | $[0, 1]$ | $[0, 1]$ | $Scm^{-2}$ | K_Tst.mod |
| KP | - | - | $[0, 1]$ | $[0, 1]$ | $Scm^{-2}$ | K_Pst.mod |
| Kv3.1 | $[0, 1]$ | $[0, 1]$ | $[0, 2]$ | $[0, 2]$ | $Scm^{-2}$ | Kv3_1.mod |
| Kv2-like | - | - | - | $[10^{-7}, 10^{-1}]$ | $Scm^{-2}$ | Kv2like.mod |
| $I_m$ | $[10^{-7}, 10^{-2}]$ | $[10^{-7}, 10^{-2}]$ | - | - | $Scm^{-2}$ | Im.mod |
| $I_m v2$ | - | $[10^{-7}, 10^{-2}]$ | - | - | $Scm^{-2}$ | Im_v2.mod |
| SK | - | - | $[10^{-7}, 10^{-1}]$ | $[10^{-7}, 10^{-1}]$ | $Scm^{-2}$ | SK.mod |
| $Ca_{HVA}$ | - | - | $[10^{-7}, 10^{-3}]$ | $[10^{-7}, 10^{-3}]$ | $Scm^{-2}$ | Ca_HVA.mod |
| $Ca_{LVA}$ | - | - | $[10^{-7}, 10^{-2}]$ | $[10^{-7}, 10^{-2}]$ | $Scm^{-2}$ | Ca_LVA.mod |
| gamma _CaDynamics | - | - | $[5 \times 10^{-4}, 0.05]$ | $[5 \times 10^{-4}, 0.05]$ | | CaDynamics.mod |
| decay _CaDynamics | - | - | $[20, 1000]$ | $[20, 1000]$ | $ms$ | CaDynamics.mod |

**Table S1**: Conductance density bounds for spiny/aspiny cells. Red and blue highlight across rows or columns represent specificity to excitatory (spiny) and inhibitory (aspiny) cells respectively. Note that none of the aspiny cell reconstructions in the Allen Cell-Types database has apical dendrite markings, thus the conductances on the apical dendrite in our models is specific to spiny cells.

| Feature Name | eFEL Name | Type | Description |
|---|---|---|---|
| **voltage_base** | voltage_base | Numeric | The average voltage during the last 10% before stimulus onset in mV |
| **steady_state_voltage** | steady_state_voltage | Numeric | The average voltage after the stimulus |
| **voltage_deflection** | voltage_deflection _vb_sse | Numeric | The voltage deflection between voltage base and average voltage during the last 10% of the stimulus duration in mV. |
| **sag_amplitude** | sag_amplitude | Numeric | The difference between the minimal voltage and average voltage during the last 10% of the stimulus duration in mV. |
| **sag_ratio** | sag_ratio1 | Numeric | The ratio between the sag amplitude and the maximal sag extend from voltage base. |
| **decay_time_constant_after_stim** | decay_time_constant _after_stim | Numeric | The decay time constant of the voltage right after the stimulus. |
| **Spikecount** | Spikecount | Numeric | Number of spikes in the trace, including outside of stimulus interval. |
| **spike frequency** | mean_frequency | Numeric | Mean frequency calculated as number of action potentials during stimulation divided by time between stimulus onset and last spike in Hz. |
| **time_to_first_spike** | time_to_first_spike | Numeric | Time to first spike in ms. |
| **AP_amplitude** | AP_amplitude_from _voltagebase | Numeric | Height at peak of action potential in mV from voltage base. Mean for all AP. |
| **AP_width** | AP_width | Numeric | Mean of width at -20 mV of AP in ms. Mean for all AP. |

| | | | |
|---|---|---|---|
| **AHP_depth** | AHP_depth | Numeric | Relative voltage values with respect to voltage_base at the first after-hyperpolarization. Mean for all AP. |
| **adaptation_index2** | adaptation_index2 | Numeric | Normalized average difference of two consecutive ISI starting from second ISI. |
| **ISI_CV** | ISI_CV | Boolean | The coefficient of variation of the ISI. |
| **depolarization_ block** | depol_block | Boolean | True if depolarization block is detected during spiking. |
| **check_AISInitiation** | check_AISInitiation | Boolean | True if time difference between same AP recorded at axon and soma is positive False otherwise |

**Table S2**: Ephys features used at different stages during model generation workflow. sag_ratio1 is not used in the optimization but used in the comparison between Nr5a1 and Rbp4 Cre-lines in Figure 6 of the main manuscript.

| Stage | Parameters | Frozen Parameters/Tolerance | eFEL Features | Stimulus |
|---|---|---|---|---|
| **Stage 0** | • $c_m$<br>• $e_{pas}$<br>• $g_{pas}$<br>• $R_a$ | – , – | • voltage_base<br>• steady_state_voltage<br>• voltage_deflection_vb_sse<br>• decay_time_constant_after_stim | All depolarizing subthreshold |
| **Stage 1** | • ḡIh | None, ±50% | eFEL Features at Stage 0+<br>• sag_amplitude | All depolarizing subthreshold + all hyperpolarizing subthreshold |
| **Stage 2** | • ḡNaT, ḡNaP<br>• ḡNaV<br>• ḡKT, ḡKP<br>• ḡKv3.1<br>• ḡKv2like<br>• ḡIm, ḡImv2<br>• ḡSK<br>• gamma/decay_ CaDynamics | None, ±50% | • voltage_base<br>• steady_state_voltage<br>• mean_frequency<br>• Spikecount<br>• time_to_first_spike<br>• AP_amplitude_ from_voltagebase<br>• AP_width<br>• AHP_depth<br>• adaptation_index2<br>• ISI_CV<br>• depol_block<br>• check_AISInitiation | ×2 spiking traces with maximal amplitude + Rheobase trace + Maximal subthreshold |

**Table S3**: Details of Optimization Workflow: We fit conductance densities (ḡ) on the primary neurites-apical, basal dendrites, soma and axon, with the assumption that these conductances are distributed uniformly across each section of the reconstructed morphology belonging to the neurite.

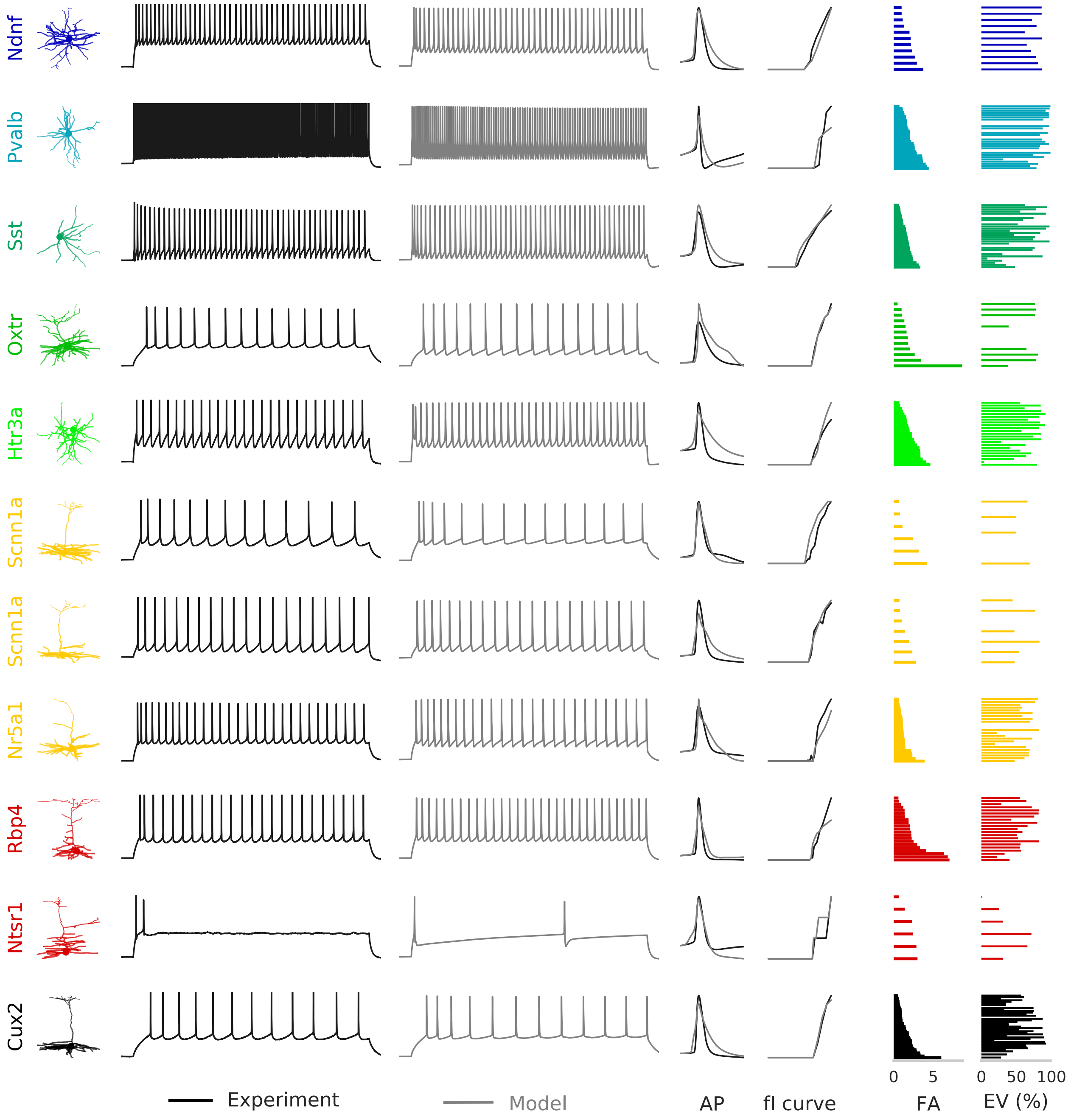


**Figure S1: Performance of representative single cell models across transgenic lines.** From left to right, Cre-line (only Cre-lines with at least 5 models are selected), morphology, experimental and model suprathreshold response, spike shape and fI curves are illustrated. The last two columns in the grid show the feature average (FA) at training and explained variance (EV) under colored noise stimulus. The representative model belongs to the cell located at the median of the feature average distribution of the member Cre-line.

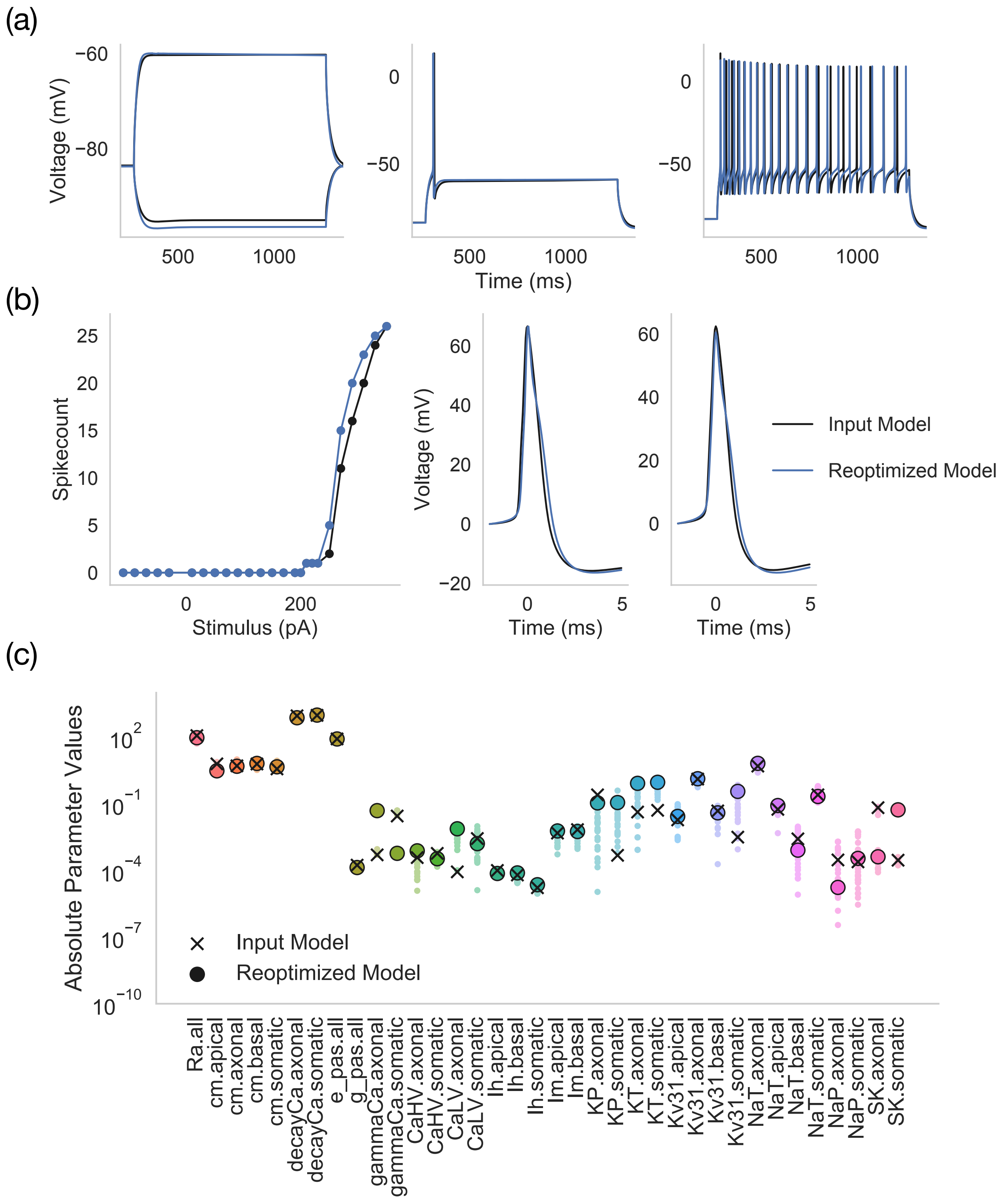


**Figure S2: Convergence of the optimization workflow for a ground truth solution.** Simulated response from the all-active model corresponding to CellID: 468193142 is fed to the workflow for a second round of optimization. **(a)** Comparison between subthreshold and suprathreshold responses as well as **(b)** fI curve and spike shapes for the input and reoptimized models**. (c)** The distribution of the hall of fame model parameters (circles) at the conclusion of the re-optimization. The best model (hof index =

0, full circle) is shown as well as the ground truth, input model (shown as ‘x’) overlaid to examine convergence. Note that the majority of the conductance parameters, especially the ones highlighted in the manuscript for downstream analysis in excitatory cells, i.e., passive parameters, h-channel conductance and axonal NaT, Kv31 have converged to their ground truth.

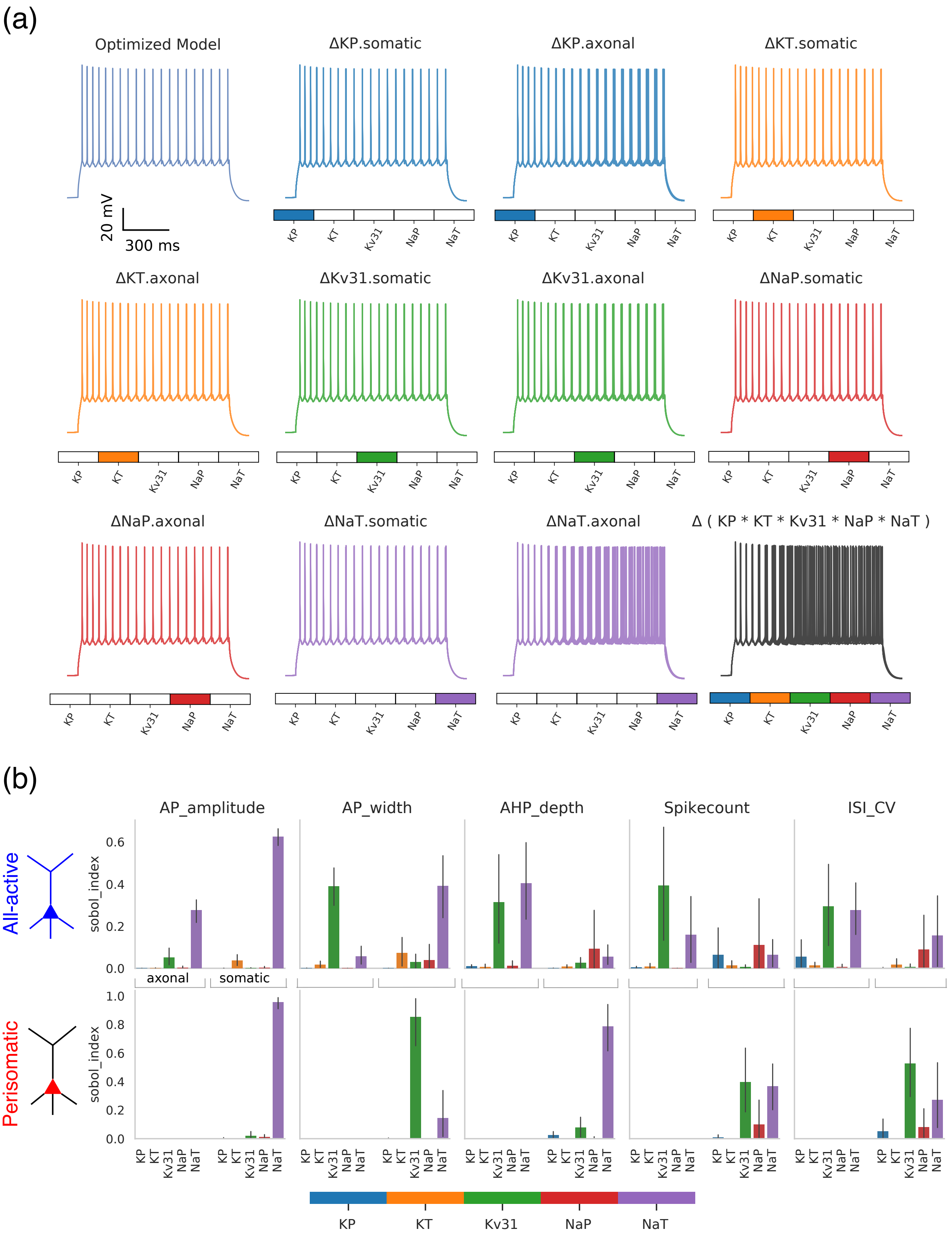
(a)
Optimized Model
20 mV
300 ms
ΔKP.somatic
ΔKP.axonal
ΔKT.somatic
ΔKT.axonal
ΔKv31.somatic
ΔKv31.axonal
ΔNaP.somatic
ΔNaP.axonal
ΔNaT.somatic
ΔNaT.axonal
Δ ( KP * KT * Kv31 * NaP * NaT )
KP
KT
Kv31
NaP
NaT
(b)
AP_amplitude
AP_width
AHP_depth
Spikecount
ISI_CV
All-active
Perisomatic
sobol_index
axonal
somatic

**Figure S3: Illustration of the sensitivity analysis and its utility in revealing mechanistic differences between all-active and perisomatic models**. **(a)** (top left) Output of the best model (least training error, i.e., hof index = 0) for Cell ID: 483101699 at 1s long stimulus of amplitude 270pA. In the following panels 100 simulations are shown for the same protocol with each of the selected $K^+$, $Na^+$ channel groups (indicated by the color bar at the bottom of each panel) individually perturbed within a ±10% tolerance level about the optimized value. Finally, at bottom right the channels are perturbed together, equivalent to the sobol analysis computations and plot the simulations for 100 such parameter combinations. Only the parameters on the color bar are modified in the illustration, the rest of the channel densities remain unperturbed from their optimized value. **(b)** The comparison between the sobol indices as it relates to spike amplitude, width, AHP depth, spike count and ISI coefficient-of-variation for all-active and perisomatic models (10 excitatory cells each, bar: mean; error bar: std). Note the absence (due to the design of perisomatic models) of axonal conductances in the bottom panel, which play a key role for the all-active spiking features (top).

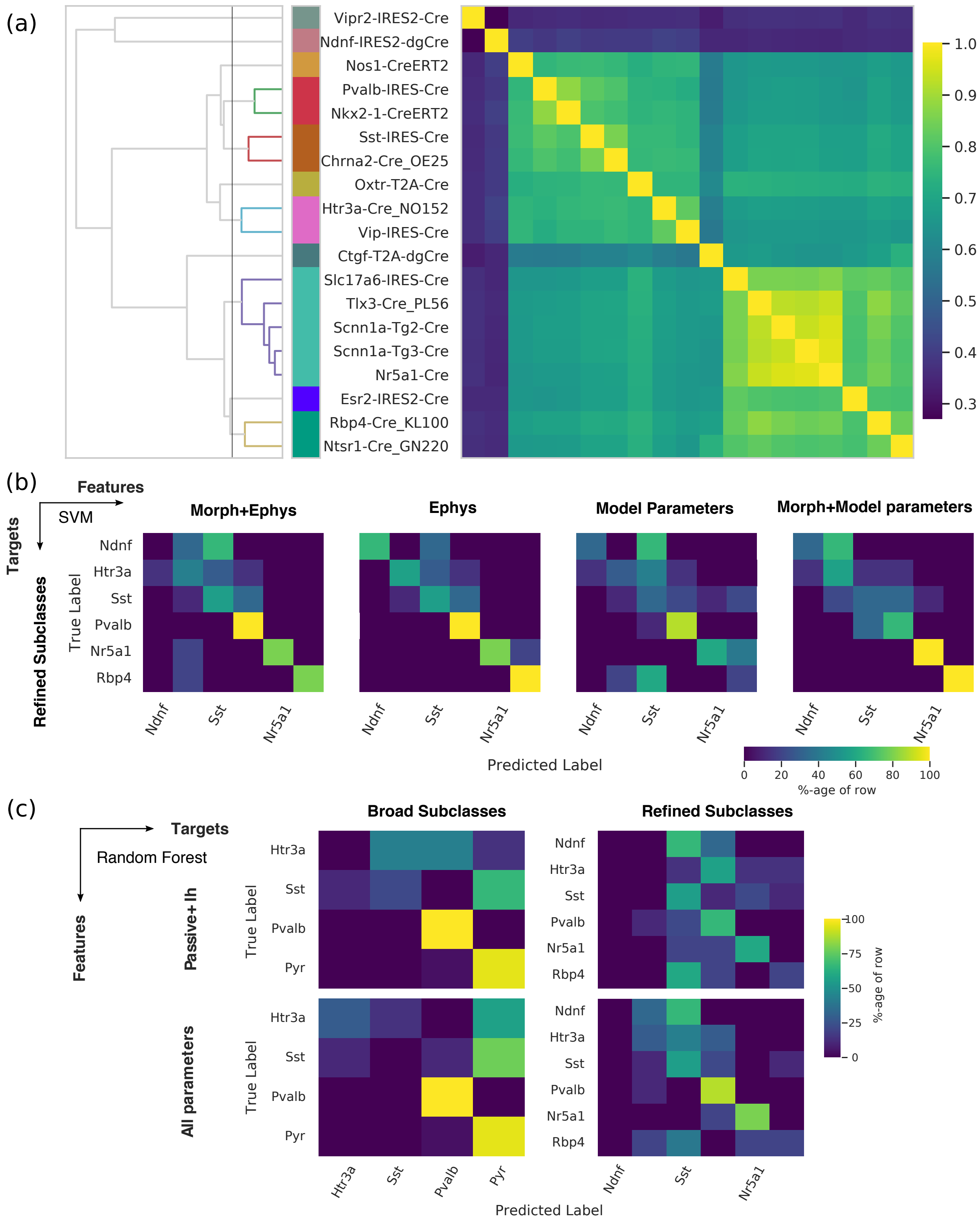


**Figure S4: Compatibility between Cre-line and transcriptomically defined cell-types and supervised classifier performance. (a)** Correlation matrix between the transcriptomic types mapped to

Cre-lines by single-cell RNA-sequencing data in a Monte Carlo fashion. The resultant Cre- to Cre-line correlation matrix (only Cre-lines from the model set are shown) is passed through hierarchical clustering resulting in similarity between the Cre-lines (left, 11 refined cell-types shown along with the dendrogram; Cre-lines of the same color are members of the same type). Confusion matrices for the **(b)** SVM classifier with 6 refined subclasses as targets and 4 feature sets (accuracy: 66%, 76%, 48% and 63% respectively) similar to **Figure 3b** and **(c)** Random forest classification (69% accuracy with both feature sets for broad subclasses; 40%, 53% respectively for reduced and all parameters respectively for refined subclasses) illustrated in **Figure 3d**.

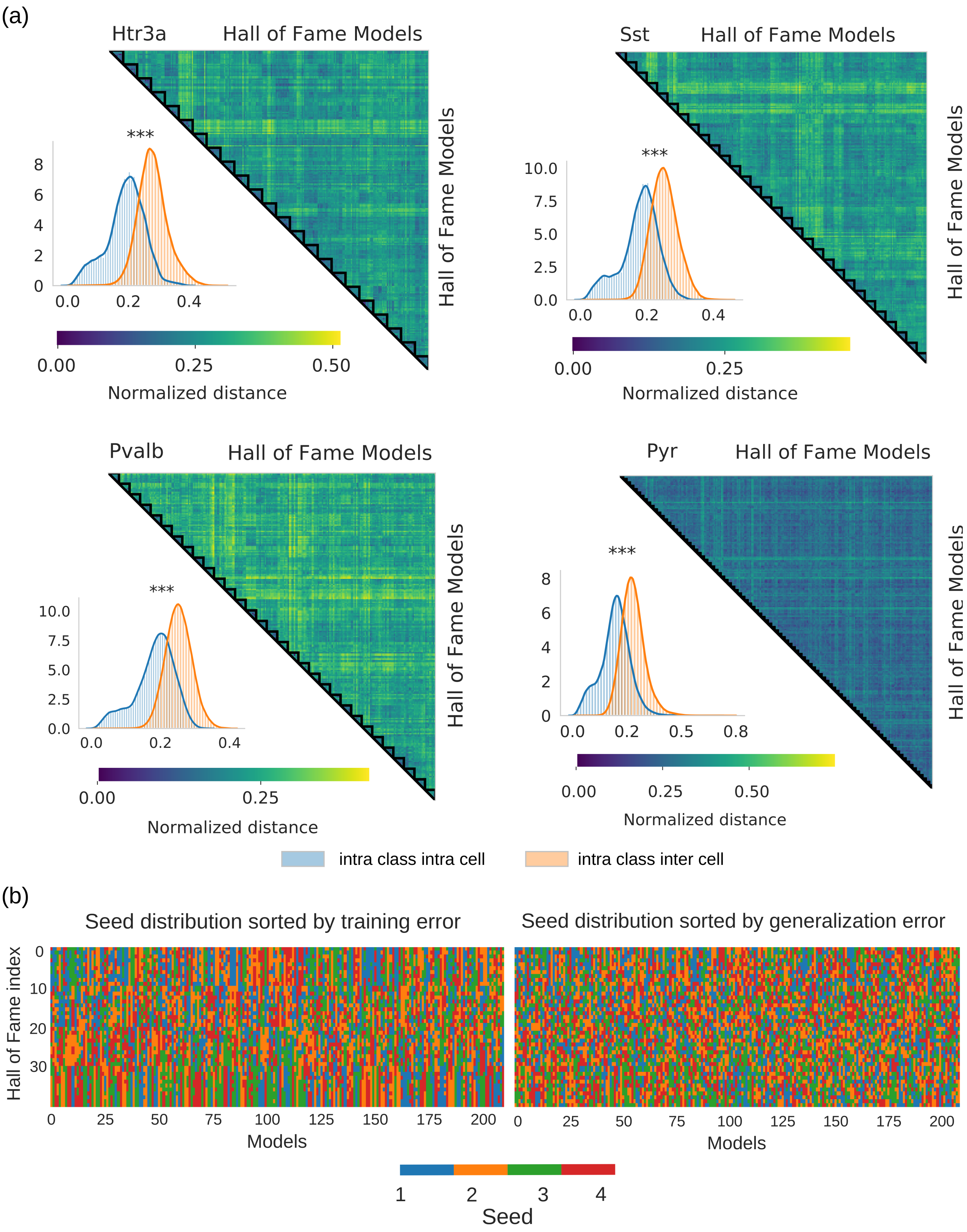


**Figure S5: Parameter diversity across cortical cell classes. (a)** Similar to **Figure 6(d)** the dispersion matrix for the hall of fame parameters within and across cell for the 4 broad subclasses. The block diagonals for 'Pyr' are compressed due to numerical advantage in comparison to the other inhibitory subclasses. For each broad subclass the intra-class intra-cell dispersion (blue) is less than intra-class inter-cell (orange). Mann-Whitney U-test; Statistical significance: *: p-val < 0.05, **: p-val $< 10^{-2}$, ***: p-val $< 10^{-3}$. **(b)** The hall of fame models for each cell is not dependent on the initialization. The single-color strands belonging to one seed when sorted to training error (left) is lost when the models are arranged according to the generalization error, i.e., performance on unseen stimulus set. This points to an existence of attraction basin for this highly constrained optimization formulation.

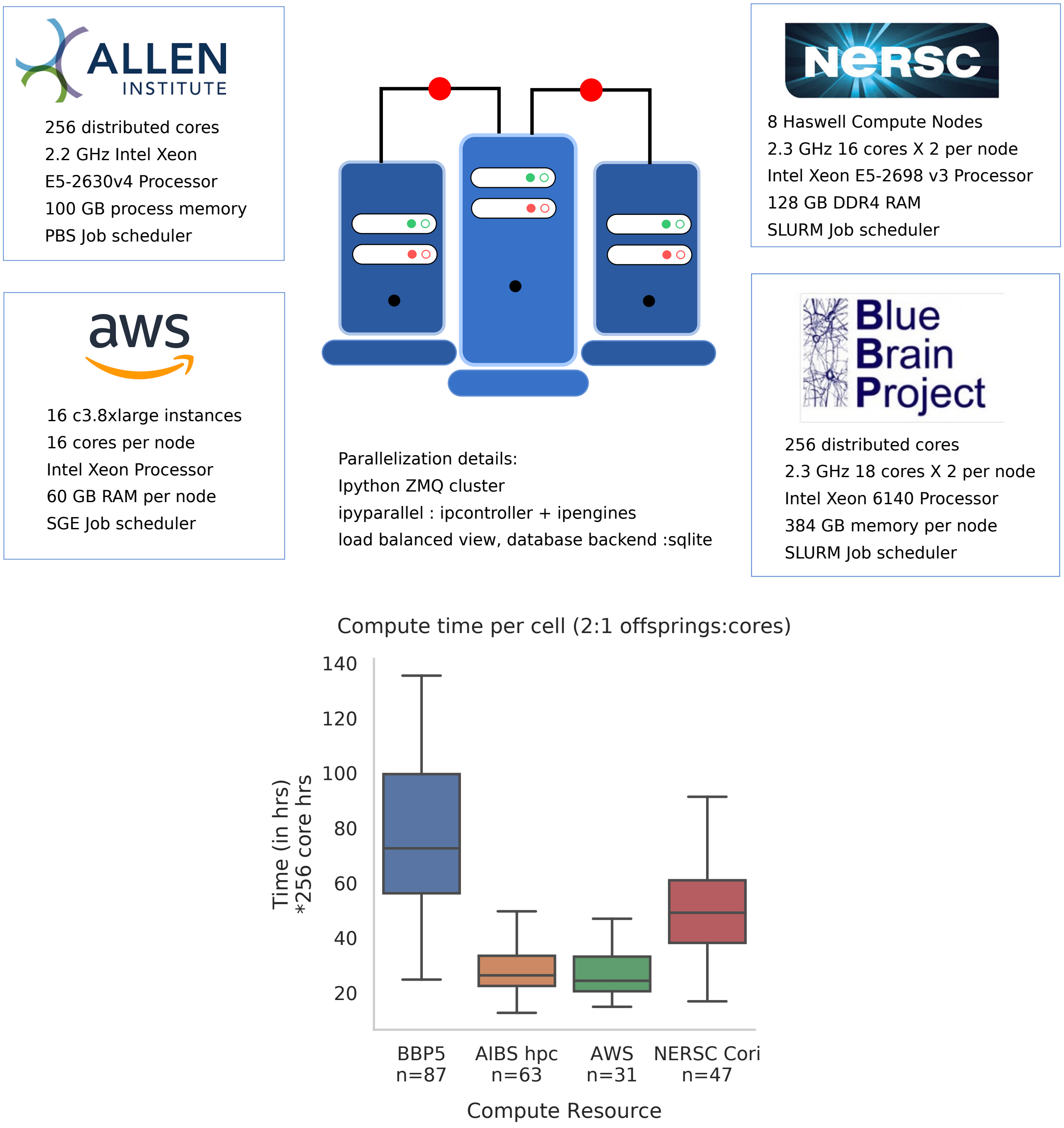


**Figure S6: Description of the computational resources employed during the model generation workflow.** Resources utilized thus far include the in house Allen Institute for Brain Science High Performance Computing (AIBS hpc) cluster, Amazon Web Services (AWS) EC2 instances in conjunction with Wasabi cloud storage as AWS S3 substitute, Cori supercomputers at National Energy Research Scientific Computing Center (NERSC) and Supercomputing infrastructure at Blue Brain Project (BBP5), École Polytechnique Fédérale de Lausanne, Switzerland. In total close to 3.5 million core hours are utilized across these four Linux based systems for the ~230 models. Reduction in compute time and subsequently core hours in AIBS hpc and AWS EC2 instances is mainly due to an algorithmic improvement in the latter stages of the model generation timeline, where solutions were discarded for which evaluation exceeded a 300s cut-off. Number of requested cores at each job is equal across machines.

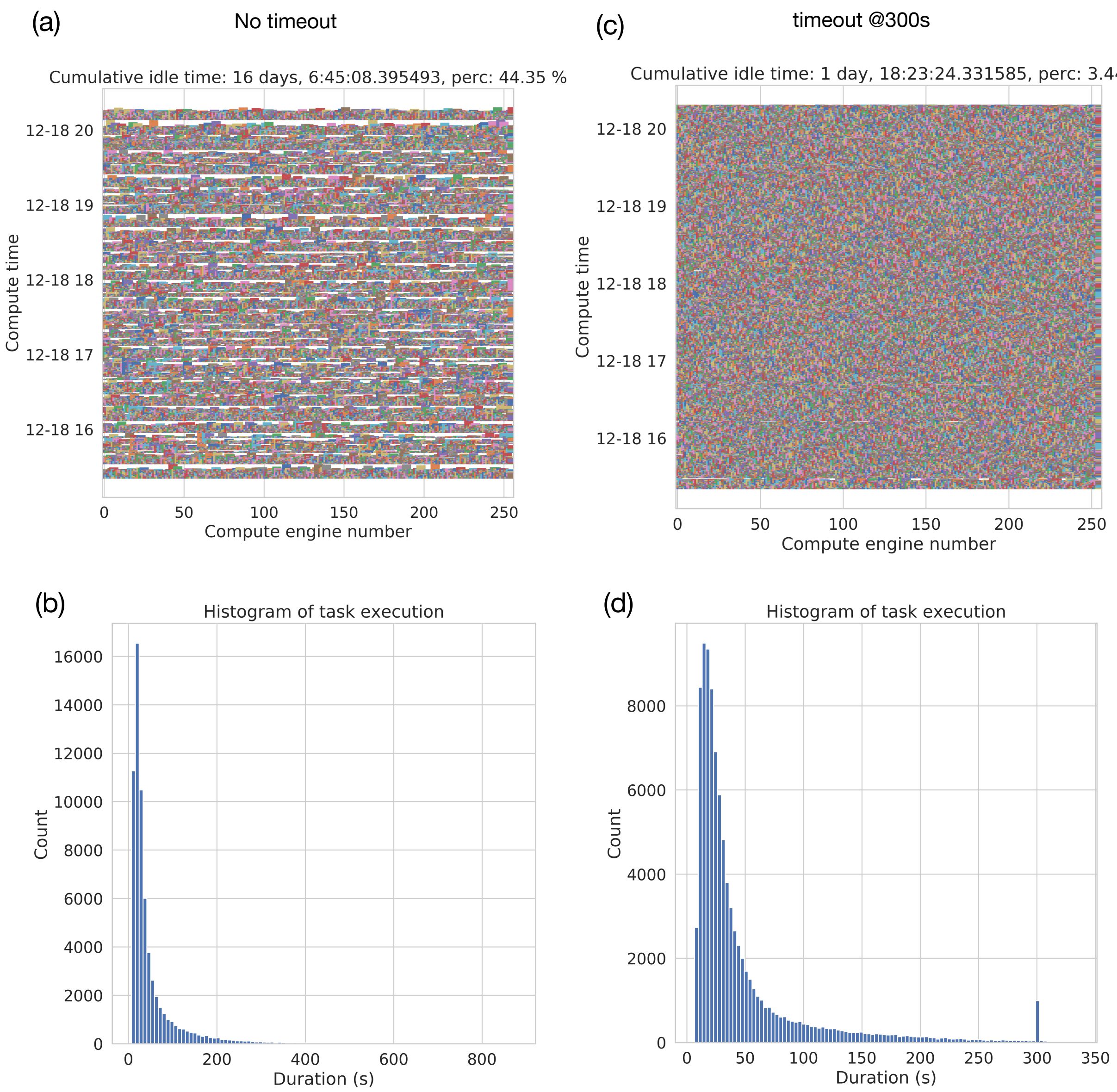


**Figure S7: Effect of adding timeout functionality within BluePyOpt**. **(a)** Without timeout the evolutionary algorithm occasionally encounters a pathological parameter combination resulting in a stiff differential equation. The blank spaces represent the idle time for the rest of engines in a distributed infrastructure. **(b)** The histogram of the task execution shows that the evaluation time for rare individuals go up to 900s. **(c)** In comparison adding a timeout of 300s enables us to run a controlled optimization in the HPC clusters with drastic reduction in idle time (44% to 3%). The jump in the histogram **(d)** at 300s means that the engines are reassigned to evaluate new individuals without completing the numerical integration when the cut-off time is hit. For this illustration we have used the stage 2 for model generation of CellID: 483101699 under identical parameter bounds.
